# Taking stock of the past: A psychometric evaluation of the Autobiographical Interview

**DOI:** 10.1101/2021.12.22.473803

**Authors:** Amber W. Lockrow, Roni Setton, Karen A.P. Spreng, Signy Sheldon, Gary R. Turner, R. Nathan Spreng

## Abstract

Autobiographical memory (AM) involves a rich phenomenological re-experiencing of a spatio-temporal event from the past, which is challenging to objectively quantify. The Autobiographical Interview (AI; Levine et *al.*, 2002, *Psychology & Aging*) is a manualized performance-based assessment designed to quantify episodic (internal) and semantic (external) features of recalled and verbally conveyed prior experiences. The AI has been widely adopted yet has not undergone a comprehensive psychometric validation. We investigated the reliability, validity, association to individual differences measures, and factor structure in healthy younger and older adults (N=352). Evidence for the AI’s reliability was strong: the subjective scoring protocol showed high inter-rater reliability and previously identified age effects were replicated. Internal consistency across timepoints was robust, suggesting stability in recollection. Central to our validation, internal AI scores were positively correlated with standard, performance-based measures of episodic memory, demonstrating convergent validity. The two-factor structure for the AI was not well-supported by confirmatory factor analysis. Adjusting internal and external detail scores for the number of words spoken (detail density) improved trait estimation of AM performance. Overall, the AI demonstrated sound psychometric properties for inquiry into the qualities of autobiographical remembering.

---

Autobiographical memory (AM) is a multifaceted form of explicit memory for personal life experiences (Conway & Pleydell-Pearce, 2000; Moscovitch et al., 2005; Rubin, 1988). AM retrieval is characterized by the phenomenological recollection, or re-experiencing, of a prior personal event including its content and spatiotemporal context. This process of re-experiencing situates aspects of AM within the broader domain of episodic memory. AM also involves accessing information from semantic memory. This information includes knowledge surrounding the event, which is not specific to a particular episode, but reflects a generalized and personal knowledge of concepts, facts and meaning (Renoult et al., 2012; Irish et al., 2013, 2012). The distinction between different types of information is necessary to measure the episodicity of recollection, yet distinguishing between episodic versus semantic features of naturalistic recollection presents unique measurement challenges.

The Autobiographical Interview (AI) was introduced by Levine and colleagues (2002) to quantify and dissociate episodic and semantic features of AM recall. This semi-structured interview elicits a verbal recounting of specific personal events, sampled across the lifespan. These narratives are transcribed and scored to characterize informational units, or details, within each memory. Details specific to the event, place, and perceptual information are identified as “internal details.” Internal details reflect qualities of the recollective experience and are considered a metric of episodic memory. Details not specific to the event, including broader conceptual and personal information that fall within the domain of semantic memory are coded as “external details,” in addition to repetitions and off-topic comments

In the first published study involving the AI, younger adults showed a bias towards reporting more internal episodic details whereas older adults showed a bias towards reporting more semantic details (Levine et al., 2002). This pattern converged with well-established age-related effects of reduced episodic memory and greater semantic knowledge in older adults (Park et al. 2001; Spreng & Turner, 2019). Additionally, the authors demonstrated convergent validity with the episodic specificity rating from the Autobiographical Incident Schedule of the Autobiographical Memory Interview (AMI), a coarser, single factor measure of AM, which rates verbal event recall on a continuum from general knowledge to episodic detail (Kopelman et al., 1989, 1990).

The AI has been widely adopted to characterize the involvement of episodic memory during AM in healthy participants and patient populations (see https://levinelab.weebly.com/ai-testing.html; Miloyan et al., 2019 for review). Consistent with the original report, healthy younger adults provide more internal details and fewer external details than older adults when recalling past personal experiences (Addis et al., 2008; Addis et al., 2010; De Brigard et al., 2016; De Brigard et al., 2017; Ford et al., 2014; Gaesser et al., 2011; Madore et al., 2014; Peters et al., 2019; Robin & Moscovitch, 2017; Spreng et al., 2018; St Jacques & Levine, 2007; Vandermorris et al., 2013; Zavagnin et al., 2016). Patients with medial temporal lobe lesions and episodic memory impairment provide fewer internal details compared to controls (Dede, Franscino et al., 2016; Dede, Wixted et al., 2016, Gilboa et al., 2006; Hilverman et al., 2016; Kirwan et al., 2008; Kwan et al., 2010; Kwan et al., 2015; Kwan et al., 2016; Miller et al., 2020; Race et al., 2011; Rosenbaum et al., 2004; Rosenbaum et al., 2008; Rosenbaum et al., 2009; Squire et al., 2010; Steinvorth et al., 2005). Individuals with Alzheimer’s disease and amnestic mild cognitive impairment, also produce fewer internal details compared with controls (Addis et al., 2009; Bastin et al., 2013; Coelho et al., 2019; Gamboz et al., 2010; Meulenbroek et al., 2010; Murphy et al., 2008; Sheldon et al., 2015). These group comparisons provide converging evidence to support the validity and reliability of the AI as a measure of AM. Some studies have gone beyond group differences to examine the individual difference properties of AM (Palombo et al., 2013; Palombo et al., 2018; Sheldon et al., 2016). Doing so has shed light on how brain structure and function shape the episodic quality of recollection on the AI (e.g., Sheldon et al., 2016; Clark et al., 2022). Yet the reliability of trait-level AM has not been formally evaluated. The AI appears to have modest test-retest reliability, with only small differences in details observed across test sessions (e.g., Barry et al., 2020). An open question is to what degree AM is consistent across memories from different time periods, in line with trait-like recollective styles, or varies as a function of remoteness.

While the AI has demonstrated internal validity and reliability, few studies have interrogated how the AI intersects with relevant psychological constructs, and those that have would benefit from further replication. Relationships to theoretically associated cognitive and psychological factors have also remained under-investigated. AM supports or is linked to an array of non-mnemonic constructs including executive function, depression symptoms, temporal discounting, social cognition, and facets of personality. Executive function supports episodic memory retrieval (Abellán-Martínez et al., 2019; Conway & Pleydell-Pearce, 2000; Yubero et al., 2011), low mood has been linked to less episodic specificity (Brittlebank et al., 1993; Hitchcock et al., 2014; Kuyken & Dalgleish, 1995; Liu et al., 2013; Williams & Broadbent, 1986; Williams & Scott, 1988; Williams et al., 2007; Wilson & Gregory, 2018), social cognition and AM depend on the same underlying system (Buckner & Carroll, 2007; Buckner et al., 2008; Gaesser, 2013; Gaesser & Schacter, 2014; DuPre et al., 2016; Rabin et al., 2010; Spreng et al., 2009; Spreng & Grady 2010; Spreng & Mar, 2012), and personality measures have been associated with autobiographical experience and expression (Adler et al., 2007; McAdams et al., 2004; Rasmussen & Berntsen, 2010; Rubin & Siegler, 2004). Yet these have not yet been evaluated in relation to the AI. AM has also been proposed as a relevant tool in assessing present and future rewards by providing a scaffold for imagining future scenarios (Gilbert & Wilson, 2007; Ersner-Hershfield et al., 2009; Bulley et al., 2016). Lempert and colleagues (2020) investigated this possibility using AI measures and reported that the ability to limit discounting of temporally distant rewards was associated with episodic autobiographical memory performance. A comprehensive psychometric evaluation of the AI, and its comparison to standard laboratory measures of memory and cognitive function, has not been undertaken.

In the present study we provide a psychometric assessment of the AI in a large sample of healthy younger and older adults. Our aims were to (i) examine the reliability and validity of the AI, (ii) assess relationships with individual difference measures of psychological and cognitive function, and (iii) test whether a two-factor solution, capturing the distinction between internal and external event details, accurately reflects the data structure. To address these aims we derived eight metrics from the AI. The first three are consistent with the original protocol (Levine et al, 2002): Number of internal details (internal count), number of external details (external count), and the ratio of internal to total details (ratio score). We also derived two additional measures, internal and external density scores, which divide detail counts by the total number of words spoken (Spreng et al., 2018). Semantic detail count and density were examined as more direct measures of semantic information. Number of words spoken was also included as a distinct variable of interest. Thus, a fourth aim was to examine whether controlling for verbosity would impact the psychometric properties of the AI.

To address these aims we conducted five sets of analyses. First, we evaluated the reliability of the AI, examining inter-rater reliability and internal consistency, which included AI scores’ stability across event memories, associations with one another, associations with participant ratings (e.g., vividness, personal relevance, emotionality, rehearsal frequency) and associations with scorer summary ratings. Next, we evaluated how AI scores varied by demographic factors across our sample, including age, gender and education level. Third, we examined convergent validity by testing associations with laboratory measures of memory and cognition. We then tested associations between the AI and factors previously implicated in AM including depression, decision-making, social cognition and personality. Finally, we conducted two sets of factor analyses. First we tested the two-factor internal/external model of the AI with confirmatory factor analysis (CFA). Follow-up exploratory factor analyses (EFA) were conducted to examine alternative factor structures.

Based on the evidence summarized above, we expected the AI to be a reliable and valid measure of AM. Specifically, we predicted high inter-rater reliability and internal consistency. Overall, we expected strong positive associations among AI scores although controlling for verbosity may alter the magnitude of these associations. We expected to replicate the commonly observed age effect of more internal details and fewer external details for younger adults compared to older adults, but had no *a priori* predictions about the impact of demographic variables. Based on previous work, we also predicted that the AI would be significantly associated with other psychological constructs. We expected that laboratory measures of episodic memory would be associated with internal measures, while laboratory measures of semantic memory would be associated with external measures. We predicted that endorsement of depression symptoms would negatively correlate with internal details and positively with external details. We predicted that greater temporal discounting, prosocial cognitive traits, and the big five personality trait openness/intellect would all be associated with higher levels of internal detail. Finally, we expected that the CFA would support the two-factor AI model. We expand on our predictions for each of the analyses in the corresponding results sections below.

## Methods

### Participants

Participants were recruited from greater Ithaca, NY, US and Toronto, Ontario, Canada and were screened to exclude psychiatric, neurological, or other illnesses that could impair cognitive functioning. 203 younger adults and 158 older adults completed all primary measures of interest. Two younger adults and four older adults were excluded for having scores below 27/30 on the Mini Mental Status Exam (Folstein, Folstein, & McHugh, 1975) combined with fluid cognition scores below a national percentile of 25% on the NIH Cognition Toolbox (Weintraub et al., 2014). Two older adults were excluded for scores above 20/30 on the Geriatric Depression Scale (Sheikh & Yesavage, 1986), indicative of moderate or severe depressive symptoms. One older adult was excluded due to extensive confabulatory statements post-experimental session. The final sample included 201 younger adults (114 women, *M*= 22.4 years; *SD*= 3.28 years; range= 18-34 years) and 151 older adults (82 women, *M* = 68.8 years; *SD=* 6.67; range= 60-92 years). Demographic information for this sample is shown in Table 1. We have recently reported on a subset of these participants with neuroimaging data (Setton et al., 2022a; Setton et al., 2022b).

**Table 1.** Participant Demographics. Gender distributions, age in years, and education level in years are reported. For the full sample, mean and standard deviation are not included for age given its bimodal distribution. W = Women, M = Men, *M* = Mean, *SD* = Standard Deviation

|  | Gender |  | Age |  | Education |  |  |  |
| --- | --- | --- | --- | --- | --- | --- | --- | --- |
|  | W | M | *M* | *SD* | Range | *M* | *SD* | Range |
| Younger Adults | 114 | 87 | 22.4 | 3.28 | 18-34 | 15.0 | 1.78 | 12-21 |
| Older Adults | 82 | 69 | 68.8 | 6.67 | 60-92 | 17.4 | 3.04 | 12-26 |
| All | 196 | 156 |  |  | 18-92 | 16.0 | 2.67 | 12-26 |

### Procedure

Participants completed the measures described below over several testing sessions as part of a comprehensive behavioral assessment protocol examining goal-directed behavior.

#### Autobiographical Interview

All participants completed the AI as specified in the original Levine et al. (2002) protocol. Older adults provided detailed descriptions of one event from five different time periods: childhood (up to age 11), teenage years (between age 11 and 18), early adulthood (between age 18 and 30), middle adulthood (between age 30 and 55), and late adulthood (within the previous year). Younger adults provided detailed descriptions of events from three periods: childhood, teenage years, and younger adulthood. For each period, participants were asked to describe an event tied to a specific time and place. Recall for the event was examined at three probe levels: free recall (uninterrupted description of the memory), general probe (general questions to elicit further details), and specific probe (specific, targeted questions to elicit details from different categories). We collapsed across free recall and general probe conditions for all metrics reported here as our focus was on spontaneous, uncued, participant-generated recollection. Note that the original study (Levine et al., 2002) reported no differences in the pattern of results for free recall and general probe. The inclusion of details from specific probe may boost the overall number of details, but is not expected to alter the pattern of results (see Levine et al., 2002). After recalling each event, participants rated the vividness, emotional change, significance (then and now), and rehearsal frequency on a 6-point Likert scale.

Participant interviews were audio recorded, anonymized, and transcribed verbatim by trained research assistants. Each transcription was then quality checked against the audio recording by a different researcher to ensure accuracy. All interviews were double-scored by two different research assistants who were trained on the protocol provided by Dr. Brian Levine and blinded to study hypotheses. Participant gender and age group may have been evident based on interview content but was not specified to transcribers or scorers. For each memory, scorers identified individual informational units, or details. Details were categorized as either “internal” or “external.” Internal details were those having to do with the identified event and were specific in time and place. External details were those that did not include information about the specified event, were not specific in time or place, or consisted of semantic information (i.e., conveyed facts that temporally extended beyond the event, world knowledge). Both internal and external details could be broken down into sub-categories of event, place, time, perceptual, and emotion/thought details. External details also included semantic, repetition, or other details. Detail counts were then tallied. As part of the scoring protocol, scorers also provided summary ratings indicating the level of detail provided. Scorers assigned an overall score for each memory based on ratings for place, time, perceptual, and emotion/thought information. Scores were also provided for time integration, episodic richness, and a global rating of detail and specificity (consistent with the episodic specificity score from the Autobiographical Incident Schedule of the Autobiographical Memory Interview (Kopelman et al., 1990): abbreviated as AMI for the purposes of this paper).

*AI Metrics.* Eight dependent variables were calculated from participant interviews:

1. Internal count
2. External count
3. Semantic count
4. Ratio score
5. Internal density
6. External density
7. Semantic density
8. Word count

Internal count (1), external count (2), and the ratio of internal to total details (4) were calculated following the procedure reported in Levine et al. (2002). To account for variation in verbosity, detail counts were divided by the word count for each memory, resulting in measures of internal (5) and external density (6; Spreng et al., 2018). To separate semantic information from non-mnemonic informational units, we also directly examined semantic count (3) and density (7). We also evaluated word count as a separate variable of interest (8). All dependent variables were calculated for each memory. Composite scores were calculated by averaging across all memories. In addition to these primary AI metrics, we also examined average self-report and scorer ratings.

### Laboratory measures of Episodic Memory, Semantic Memory, and Executive Function

Index scores of episodic memory, semantic memory, and executive function were derived from the NIH Cognition Toolbox and in-lab tasks. Measures of episodic memory included: Verbal Paired Associates (Wechsler, 2008), the Associative Recall Paradigm (Brainerd & Pressley, 2013), the NIH Rey Auditory Verbal Learning Test (Weintraub et al., 2014), and the NIH Picture Sequence Memory task. Measures of semantic memory included: the Shipley-2 Test of Vocabulary (Shipley et al., 2009), the NIH Reading Recognition Task, and the NIH Picture Vocabulary Task. Measures of executive function included: NIH List Sorting task, the NIH Card Change Sort Task, the NIH Flanker Task, a Reading Span task (Daneman & Carpenter, 1980), and the Trail Making Task (reaction time for section B minus reaction time for section A) (Reitan, 1958). Index scores were calculated by z-scoring each measure and averaging across measures within each domain. Two younger participants’ semantic index scores were winsorized for outlying performance.

A subsample of 148 younger adults and 90 older adults had complete data for the Remember/Know paradigm (R/K; Tulving, 1972; Tulving 1985). Participants were excluded if they provided Remember responses for more than 90% of recognized items, which limited the number of familiarity trials (e.g., Stamenova et al., 2017). Fifty-two participants were excluded on this basis: 23 younger adults and 29 older adults. Recollection and familiarity scores were derived according to standardized methods (Söderlund et al., 2008).

### Measures of depression, decision-making, social cognition and personality

A subsample of 67 younger adults and 93 older adults completed a measure of temporal discounting (Loeckenhoff et al., 2011). Temporal discounting was evaluated with a computerized forced choice task in which participants selected whether they would prefer to receive small magnitude rewards now or at varied dates in the future (7-180 days). Dependent variables included area under the curve (Myerson et al., 2001), reward index (Boettiger et al., 2007), and the proportion of patient (delayed) choices. To facilitate comparisons with prior work associating the AI with temporal discounting (Lempert, MacNear, et al., 2020), we computed an additional “perceptual/gist ratio” (computed as [(internal time count + internal place count + internal perceptual count) / total internal count]) from the AI scores.

A subset of participants completed additional online self-report questionnaires on Qualtrics. Fifteen younger adults and three older adults were excluded based on failed attention checks. Thus, the maximum sample for analyses that included self-report questionnaires was 162 younger adults and 125 older adults. Depressive symptoms were measured with the Beck Depression Inventory (Beck et al., 1996) for younger adults and the Geriatric Depression Scale (Sheikh & Yesavage, 1986) for older adults. Social Cognition measures included the Interpersonal Reactivity Index (Davis, 1983), the Toronto Empathy Questionnaire (Spreng et al., 2009), and the Reading the Mind in the Eyes Task (Baron-Cohen et al., 1997). Personality was evaluated with the Big Five Aspect Scales (DeYoung et al., 2007).

### Analysis

All analyses were conducted with the R statistical software version 4.0.0 (R Core Team, 2020). The packages used included corrplot (Wei & Simko, 2017), ggplot2 (Villanueva & Chen, 2019), lme4 (Bates et al., 2014), lmerTest (Kuznetsova et al., 2017), lavaan 0.6-7 (Rosseel, 2012), and psych (Revelle, 2012). In order to maximize statistical power, our primary analyses leveraged the entire sample of participants where appropriate, controlling for age group and gender. We also conducted parallel analyses within the two age groups controlling for gender. Results are summarized in the main text and depicted in Supplementary Figures.

Potential outliers on AI metrics were evaluated using the median absolute deviation method (Leys et al., 2013). In total, 121 observations were winsorized prior to analysis. 37 participants’ data contained one outlier (14 younger adults and 23 older adults), and 18 had more than one (9 younger adults and 9 older adults). Outlier correction did not change the pattern of results reported here.

Throughout the manuscript, associations were examined among AI metrics (e.g. internal and external count) and between specific AI metrics and other measures with hypothesis-driven relationships (e.g., internal count and episodic memory). For completeness and transparency, relationships between potentially unpredicted associations are also reported. In order to assess validity and to guide future work, uncorrected p-values are reported (see Rothman, 1990; Saville, 1990; Althouse, 2016). Predicted associations are indicated in both the figures and the main text. Significant non-predicted associations are shown in figures and additionally flagged if they survived Bonferroni correction (α = .05) based on the number of unpredicted correlations conducted within each set of analyses. All correlation magnitudes are reported in figures to provide estimation of effect sizes based upon the current sample.

#### (i) Reliability of the Autobiographical Interview

##### Inter-rater reliability

Scoring of autobiographical events is manualized and involves significant training to set reliability criteria. Scorers must identify the primary event in the narrative, demarcate details, and determine whether these details are internal or external to the event. This decision process is subjective. To evaluate the AI’s inter-rater reliability, intraclass correlations were computed based on a mean-rating (*k*=2), absolute agreement, one-way random effects model (Shrout & Fleiss, 1979) between scorers. This was done for composite scores of total detail count, internal count, and external count, and for each of the sub-categories. Correlations were computed on the full sample and separately within younger and older adults. Detail counts were averaged between scorers for all subsequent analyses.

To evaluate the AI’s internal consistency, we tested the stability of detail generation across time points with all eight AI variables and subsequently examined the association between the eight AI composite scores averaged across events.

##### Internal consistency (across timepoints)

AMs are sampled from multiple events over the lifespan. AI scores are often derived by taking the average of internal and external details from multiple life events, even though remoteness has profound effects on memory (e.g. Linton, 1975; Rubin & Wenzel, 1996; Wagenaar, 1986). We first examined how detail recollection differed across time periods. We tested for detail differences between the most recent (proximal) and remote (childhood) memories with pairwise t-tests. Then, repeated-measure ANOVAs within age groups were conducted to test for detail differences across all three memories in younger adults and all five memories in older adults. Post-hoc Bonferroni corrected pairwise t-tests (α = .05) were conducted between all pairs of events and AI measures. Next, we quantified how similarly details were recalled across time periods with partial product-moment correlations (pr) controlling for age group and gender. Specifically, we computed correlations between 1) the most remote and the most recent event; 2) each event and the average of all events (akin to a modified item-total correlation); 3) each event and the average of the most recent and remote events; and 4) the average of all events and the average of the first and last events. Correlations were conducted for each of the eight AI measures of interest. As younger and older adults described a different number of memories, the full sample, controlling for age group and gender, was used for correlations between the most recent and remote events, and separate age groups, controlling for gender, for the remainder.

##### Internal consistency (AI metrics)

We next examined associations among the eight AI composite measures (i.e., averaged across all memories). Partial product-moment correlations were computed between the AI measures across all participants controlling for age group and gender. Analyses were also repeated within age groups (see Supplemental Material).

Recent work suggests that, within a single event, the amount of internal detail recalled is negatively related to the amount of external detail (Devitt et al., 2017). Given this possibility, we conducted a hierarchical linear model analysis to determine whether internal count predicted external count. An internal count by age group interaction was modeled as a fixed effect, with the memories from different time periods as intercepts and participants as random effects. We compared different model structures to determine the best fitting model (Judd et al., 2012). Our first model included random intercepts for memory and participant. Our second model also allowed internal count to vary by event. As a chi-square test indicated that fit did not differ between the models (*χ2*(2) = .24, *p* = .89), we report results from the simpler random intercept model.

##### Internal consistency (AI ratings)

In our final assessment of internal consistency, we examined how the eight primary, performance-based AI measures were related to self-reported participant ratings and scorer summary ratings. Self-report ratings were participant-reported event vividness, emotional change, significance now, significance then, and rehearsal frequency. Scorer summary ratings were from an overall score assigned by the scorer for each memory based on the level of detail included. Ratings were assigned for place, time, perceptual, emotion/thought, AMI episodic specificity, episodic richness (internal) and time integration (external). External details are only considered for the time integration rating. Since ratings were made on a Likert scale, associations were examined with partial Spearman correlations controlling for age and gender.

#### (ii) Demographic Associations with the Autobiographical Interview

##### Autobiographical Interview Performance by Gender, Age and Education

To determine gender and age differences, 2-way Analyses of Variance (ANOVAs) were conducted between gender and age groups for all eight AI measures of interest. Post-hoc pairwise t-tests with Bonferroni correction were run (α = .05) to examine pairwise differences between groups. We also conducted partial product-moment correlations to examine the relationship between education and the eight AI measures across all participants, controlling for gender and age. Correlations were repeated within age groups controlling for gender.

#### (iii) Convergent Validity of the Autobiographical Interview

Central to validation of the AI is determining whether it shares variance with other validated measures of related constructs. To evaluate convergent validity of the AI, we conducted partial product-moment correlations between the AI measures and index scores of episodic memory, semantic memory, and executive function. We additionally examined associations between the AI measures and recollection and familiarity on the R/K task. To maximize statistical power, our primary analyses leveraged the entire sample of participants, controlling for age and gender. Complementary analyses in separate groups of younger and older adults were performed, controlling for gender (see Supplementary Material).

#### (iv) Associations with Depression, Decision-making, Social Cognition and Personality

Additional associations were explored between the AI and non-mnemonic constructs that have been related to AM abilities in past studies. Partial product-moment correlations were conducted between the eight AI measures and: depressive symptoms, temporal discounting, and measures of social cognition and personality, controlling for age and gender. Analyses were conducted in the full sample with the exception of depression, which was measured with different questionnaires in younger and older adults. Repeat analyses within each age group controlling for gender can be found in Supplementary Material.

#### (v) Factor Analysis of the Autobiographical Interview

##### Confirmatory Factor Analysis

A primary feature of declarative memory is the distinction between episodic and semantic memory (Tulving, 1972). This feature is embedded within the internal and external distinction of the AI. We tested this two-factor model of the AI with a confirmatory factor analysis (CFA), which divided all of the AI sub-categories into latent variables of internal and external details. We estimated two models across participants (with age group embedded within the model): one with detail count and the second with detail density. Since some categories of details are reported more than others, which results in a skewed data distribution, we transformed the data by taking the cubed root of all values. CFAs were conducted with maximum likelihood estimation. Latent factors were standardized for free estimation of factor loadings. Model fit was evaluated with the Comparative Fit Index (CFI), Tucker-Lewis Index (TLI), and Root Mean Square Error of Approximation (RMSEA). The hypothetical CFA model structure is depicted in Figure 1. We also examined model fit as a function of count or density values. To do so, we compared the closeness of our two models to the data’s true structure using a Vuong closeness test for non-nested models (Vuong, 1989). Finally, as external details include both mnemonic and non-mnemonic information, we conducted another CFA with the same procedure but including only semantic details instead of all external details to determine if this adjusted model would demonstrate acceptable fit.

**Figure 1.**
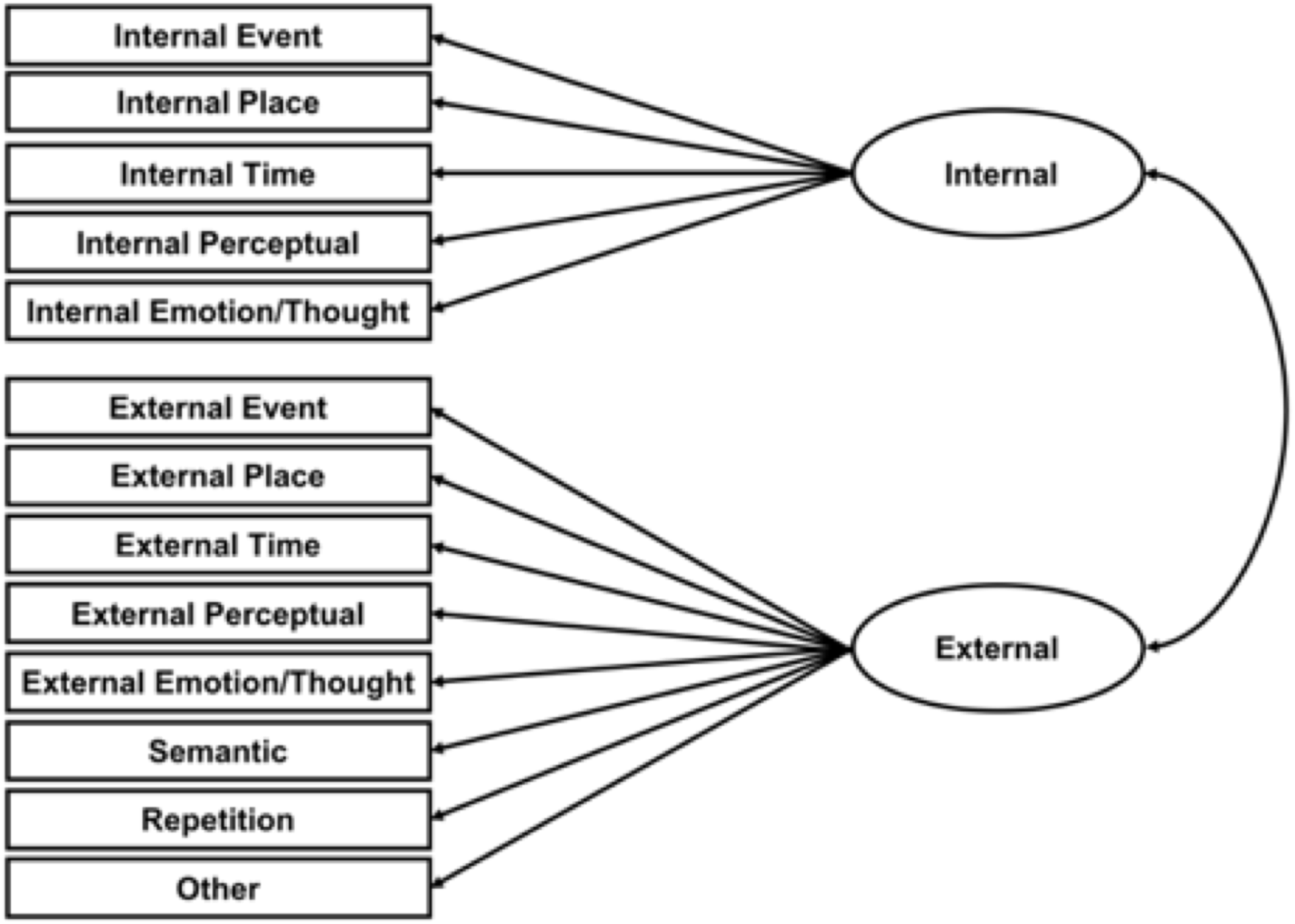
Confirmatory Factor Analysis of Internal and External Details: Hypothetical Model. The ellipses represent the latent variables of internal and external details whereas the rectangles represent observed sub-categories of detail types which fall into each latent variable category, as represented by the straight arrows. The curved arrows represent the correlation between the two latent variables.

##### Exploratory Factor Analysis

As a follow-up to the CFA, we conducted EFA to interrogate the data-driven factor structure that emerges from detail sub-categories on the AI. We conducted EFAs with varimax rotation on (i) cubed root detail counts and (ii) cubed root density scores across all participants, within younger adults, and within older adults, resulting in a total of 6 EFAs. Parallel analysis (Horn, 1965) was used to determine the optimal number of factors for each analysis.

## Results

### (i) Reliability of the Autobiographical Interview

#### Inter-Rater Reliability

Based upon the rigorous standardized protocol for training scorers, and previously reported inter-rater reliability rates (.88-.96 for internal details and .79-.96 for external details; Addis et al., 2008; Addis et al., 2010; Cole et al., 2012; Devitt & Schacter, 2020; Gaesser et al., 2011; Levine et al., 2002; Robin & Moscovitch, 2017; Terrett et al., 2016), we predicted high inter-rater reliability. Indeed, inter-rater reliability of the AI was high for internal and external details. Reliability of the average detail counts, as estimated using an intraclass correlation mean-rating model, was high across the entire sample (total r(351)=.94, p<.001; internal r(351)=.95, p<.001; external r(351)=.86, p<.001), within younger adults (total r(200)=.96, p<.001; internal r(200)=.95, p<.001; external r(200)=.88, p<.001), and within older adults (total r(150)=.92, p<.001; internal r(150)=.95, p<.001; external r(150)=.83, p<.001). Reliability for sub-categories was more variable in the full sample (r(351)=.57 – .93, p<.001), within younger adults (r(200) =.42 – .93, p<.001), and within older adults (r(150)=.44 – .94, p<.001). Individual sub-category mean-rating model correlations are provided in Table 2.

**Table 2.** Inter-Rater Reliability. Intraclass correlations estimating reliability of detail counts when averaged between raters using a mean-rating model. Correlations are reported for all participants, younger adults, and older adults for summary and sub-category detail counts. n.s. not significant; *p<.05; **p < .01; ***p<.001.

|  |  | All | Younger<br>Adults | Older<br>Adults |
| --- | --- | --- | --- | --- |
| Internal and External | Total | .94*** | .96*** | .92*** |
| Internal | Total | .95*** | .95*** | .95*** |
|  | Event | .93*** | .93*** | .92*** |
|  | Place | .74*** | .73*** | .76*** |
|  | Time | .89*** | .89*** | .88*** |
|  | Perceptual | .89*** | .86*** | .94*** |
|  | Emotion/Thought | .91*** | .91*** | .92*** |
| External | Total | .86*** | .88*** | .83*** |
|  | Event | .60*** | .76*** | .50*** |
|  | Place | .66*** | .58*** | .64*** |
|  | Time | .64*** | .48*** | .71*** |
|  | Perceptual | .70*** | .42*** | .78*** |
|  | Emotion/Thought | .60*** | .77*** | .44*** |
|  | Semantic | .85*** | .81*** | .83*** |
|  | Repetition | .59*** | .58*** | .61*** |
|  | Other | .57*** | .68*** | .45*** |

#### Internal consistency (across timepoints)

Event remoteness can have a significant impact on memory. Because the AI samples discrete events that vary as a function of remoteness, we assessed the internal consistency of the AI across timepoints in two ways.

First, we examined mean differences in each of the eight AI variables between the most recent and most remote events across all participants. This informed the stability of detail recollection with temporal distance. We predicted that remote events would have fewer details overall. Compared with remote events, recent events had higher internal count (t(351) = 10.71, p <.001, Cohen’s d =0.57), higher internal density (t(351) = 2.06, p <.05, Cohen’s d =0.11), higher external count (t(351) = 6.30, p< .001, Cohen’s d =0.34) and higher semantic count (t(351) = 4.69, p <.001, Cohen’s d =0.25). Word count was also higher for recent versus remote events (t(351) = 10.84, p <.001, Cohen’s d =0.58). These differences are depicted in Figure 2.

**Figure 2.**
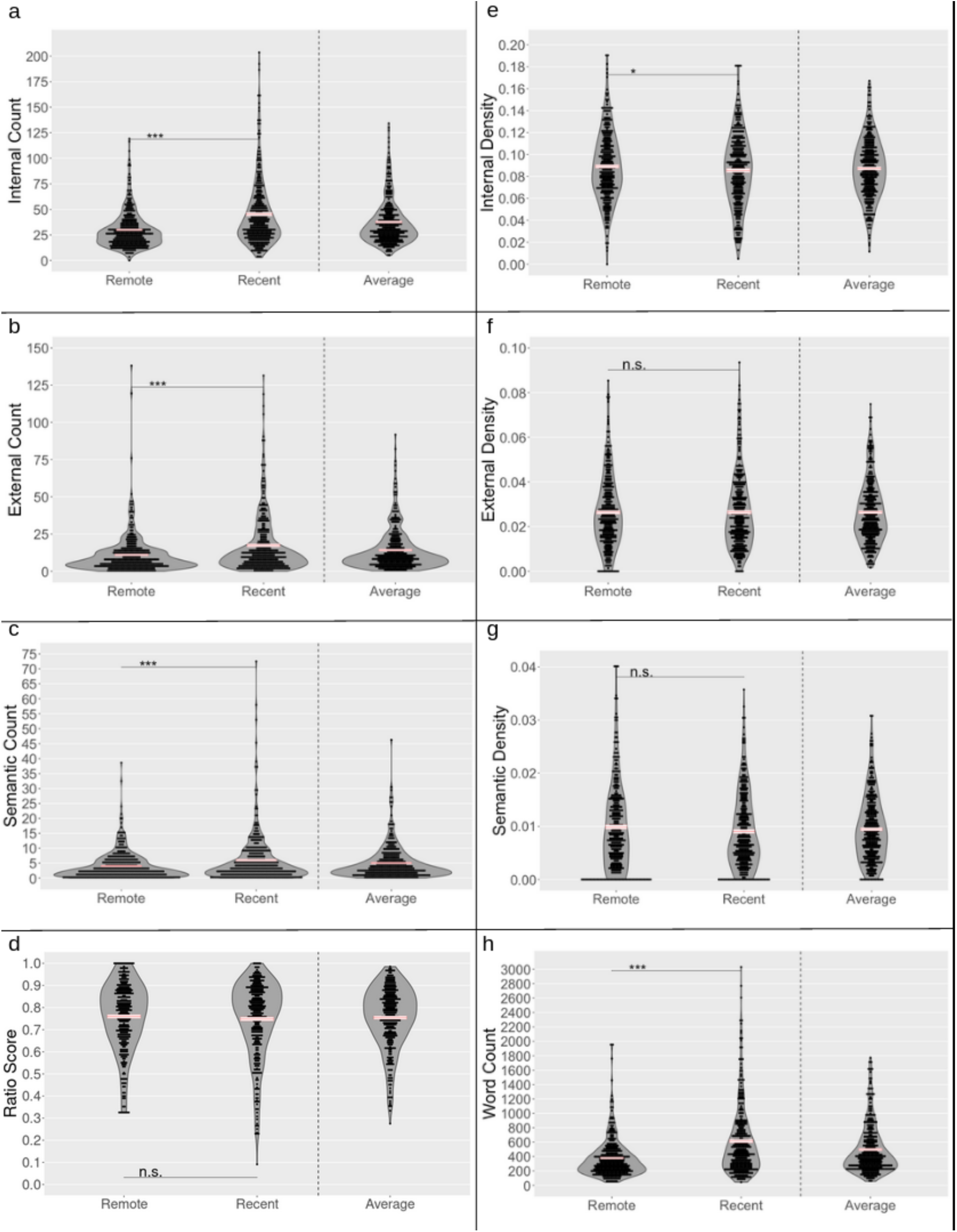
Between Memory Consistency for All Participants. The a) internal, b) external, and c) semantic counts, d) the ratio score, the density scores for e) internal f) external and g) semantic details, and h) the word count averaged across raters for the most remote and most recent memories of all participants. The averages across all memories are presented for comparison but excluded from analyses. Error bars represent the mean +/- standard error. Significant differences between recent and remote memories were identified using student’s paired t-test; n.s. not significant; *p<.05; **p < .01; ***p<.001.

Within younger adults, we tested for mean differences on all AI variables across childhood, teenage years, and recent early adulthood memories (Figure 3). We found a stepwise increase in internal count with recency (F(2, 400)=35.33, p<.001, Cohen’s f=0.42; Figure 3a). Remote childhood memories also had lower external count (F(2, 400)=13.13, p<.001, Cohen’s f =0.26; Figure 3b) and semantic count (F(2, 400)=5.20, p<.01, Cohen’s f =0.16; Figure 3c). A stepwise increase in word count for recency was also observed (F(2, 400)=38.91, p<.001, Cohen’s f =0.44; Figure 3h). No differences were observed for density or ratio scores (Fs < 2, p’s > .15; Figures 3d and 3e).

**Figure 3.**
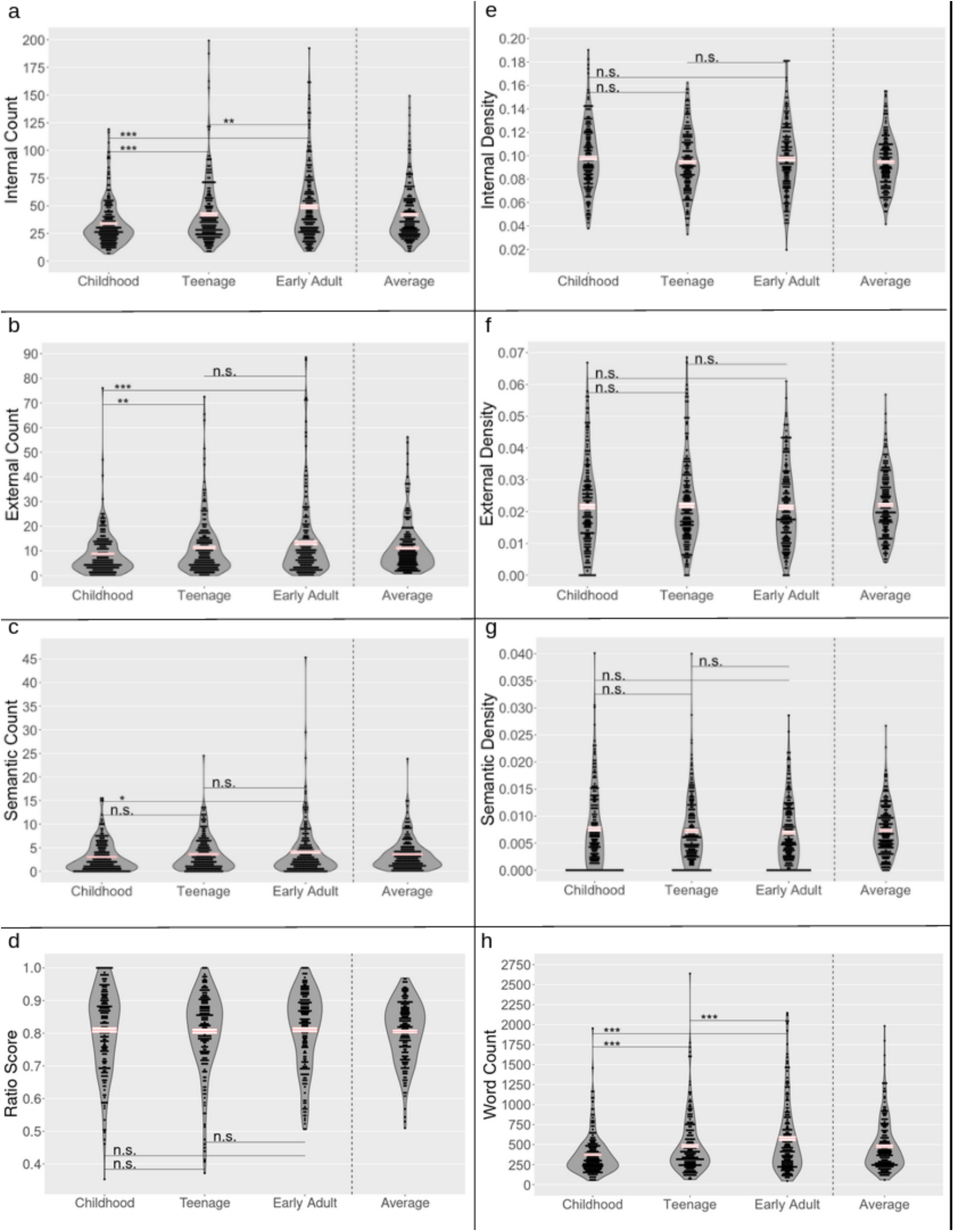
Between Memory Consistency for Younger Adults. The a) internal, b) external, and c) semantic counts, d) the ratio score, the density scores for e) internal f) external and g) semantic details, and h) the word count averaged across raters for childhood, teenage, and early adult memories in younger adults. The averages across all memories are presented for comparison but excluded from analyses. Error bars represent the mean +/- standard error. Significant differences were identified using Bonferroni corrected pairwise t-tests; n.s. not significant; *p<.05; **p < .01; ***p<.001.

Within older adults, we tested for mean differences on all AI variables across childhood, teenage years, early adulthood, middle age, and within the last year (Figure 4). Significant differences between the five events were found, with a pattern of lower values for remote events in internal count (F(4,600)=20.95,p<.001, Cohen’s f =0.37; Figure 4a), external count (F(4,600)=8.71,p<.001, Cohen’s f =0.24; Figure 4b), and semantic count (F(4,600)=6.15,p<.001, Cohen’s f =0.20; Figure 4c). Lower word count was observed in more remote events (F(4,600)=27.14,p<.001, Cohen’s f =0.43; Figure 4h). Differences were also observed for internal density (F(4,600)=4.05,p<.01, Cohen’s f =0.16; Figure 4e) and ratio score (F(4,600)=2.84, p<.05, Cohen’s f =.14; Figure 4e), with higher ratio scores and internal densities for childhood memories relative to the teenage memories. Semantic density also differed (F(4,600)=8.17,p<.001).

**Figure 4.**
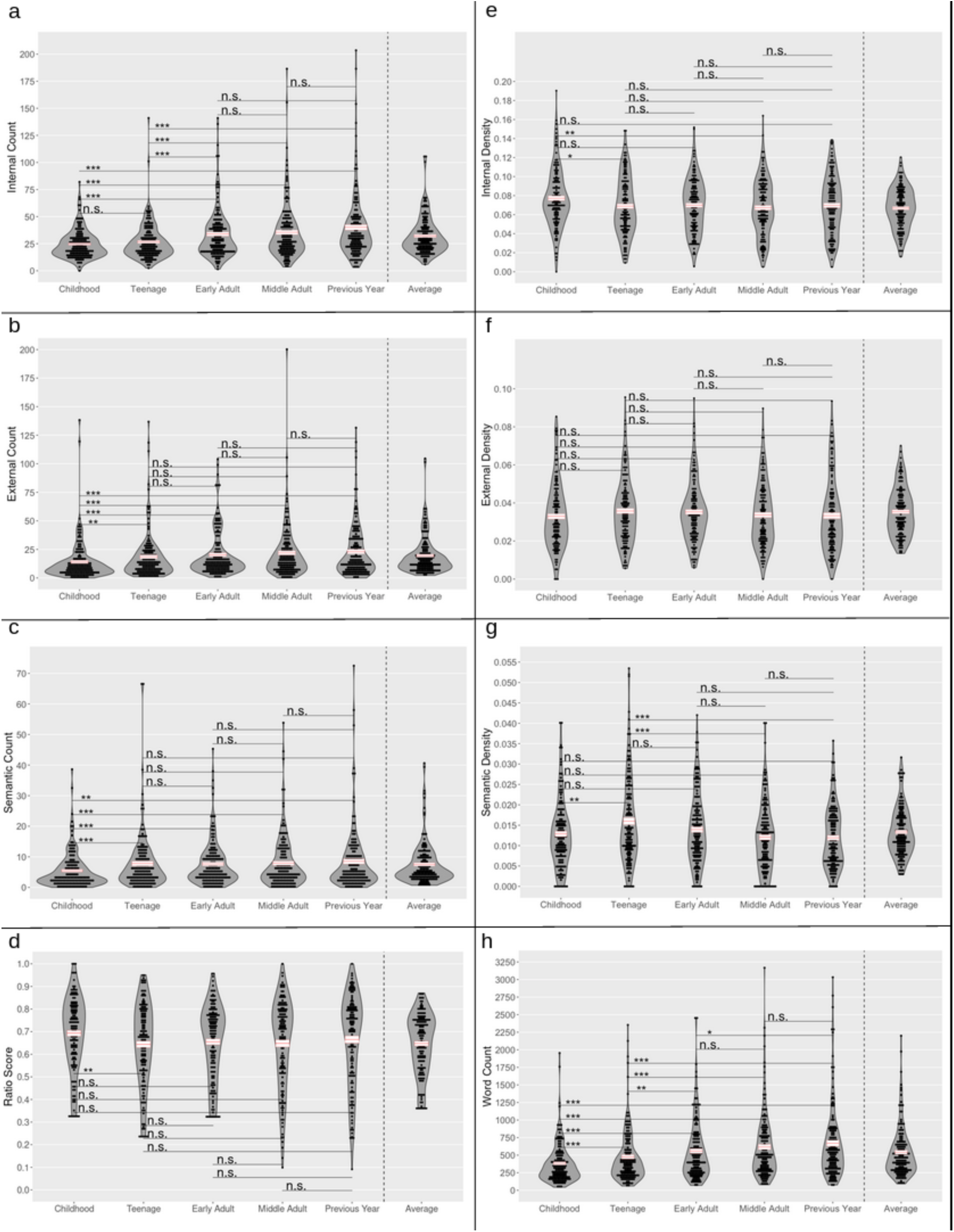
Between Memory Consistency for Older Adults. The a) internal, b) external, and c) semantic counts, d) the ratio score, the density scores for e) internal f) external and g) semantic details, and h) the word count averaged across raters for childhood, teenage, early adult, middle adult, and previous year memories in older adults. The averages across all memories are presented for comparison but excluded from analyses. Error bars represent the mean +/- standard error. Significant differences were identified using Bonferroni corrected pairwise t-tests; n.s. not significant; *p<.05; **p < .01; ***p<.001.

Recollection was robustly correlated across time periods. Recent and remote events were highly correlated across the entire sample (pr(348) = .26 – .55, p < .001), in younger adults (pr(198) = .30 – .60, p < .001), and in older adults (pr(148) = .19 – .49, p < .05). Correlations between each individual event and the average score of all events (akin to an item-total correlation) were observed across the entire sample (pr(348) = .62 - .84, p < .001), in younger adults (pr(198) = .63 – .91, p < .001), and in older adults (pr(148) = .54 – .90, p < .001). Correlations between each individual event and the average of the most recent and most remote event were also observed for the entire sample (pr(348) = .72 – .94, p < .001). This association was significant but highly variable for both the younger (pr(198) = .19-.95, p < .01) and older groups (pr(148) = .35 – .96, p < .001). Finally, we assessed the similarity of two composite measures: the average of all events and an average of the most recent and remote events. These correlations were highly positive across the entire sample (pr(348) = .81 – .92, p < .001), in younger adults (pr(198) = .83 – .95, p < .001), and in older adults (pr(148) = .77 – .89, p<.001). Correlations between event-level and composite AI measures are shown in Table 3. The duration of time between the most recent event and most remote event substantially differs between younger and older adults. However, controlling for the distance between the two events, when examining correlations between recent and remote events across the full sample, did not change the pattern of significant associations.

**Table 3.** Between Memory Consistency of Events and Composite Measures. Partial product-moment correlations of the AI between events and composite measures. Correlations are reported for all participants controlling for age group and gender, and within each age group controlling for gender. n.s. not significant; *p<.05; **p < .01; ***p<.001.

| All Participants | Internal Count | Internal Density | Proportion Score | External Count | Semantic Count | External Density | Semantic Density | Word Count |
| --- | --- | --- | --- | --- | --- | --- | --- | --- |
| Remote-Recent | .55*** | .36*** | .35*** | .37*** | .38*** | .35*** | .26*** | .55*** |
| Remote-Average | .79*** | .65*** | .64*** | .78*** | .75*** | .63*** | .66*** | .80*** |
| Recent-Average | .84*** | .80*** | .76*** | .75*** | .76*** | .74*** | .62*** | .83*** |
| Remote-Remote/Recent | .80*** | .82*** | .80*** | .75*** | .72*** | .82*** | .83*** | .79*** |
| Recent-Remote/Recent | .94*** | .82*** | .84*** | .89*** | .92*** | .83*** | .76*** | .94*** |
| Average-Remote/Recent | .92*** | .88*** | .86*** | .91*** | .89*** | .84*** | .81*** | .92*** |
| Younger Adults | Internal Count | Internal Density | Proportion Score | External Count | Semantic Count | External Density | Semantic Density | Word Count |
| Childhood-Early Adult (Remote-Recent) | .59*** | .50*** | .37*** | .47*** | .40*** | .34*** | .30*** | .60*** |
| Childhood-Average | .82*** | .74*** | .67*** | .75*** | .70*** | .63*** | .69*** | .83*** |
| Teenage-Average | .89*** | .76*** | .70*** | .82*** | .77*** | .69*** | .64*** | .91*** |
| Early Adult-Average | .86*** | .84*** | .76*** | .87*** | .87*** | .74*** | .67*** | .88*** |
| Childhood-Remote/Recent | .83*** | .86*** | .84*** | .76*** | .74*** | .84*** | .85*** | .83*** |
| Teenage-Remote/Recent | .69*** | .49*** | .29*** | .60*** | .55*** | .24*** | .19** | .75*** |
| Early Adult-Remote/Recent | .94*** | .87*** | .82*** | .93*** | .91*** | .80*** | .76*** | .95*** |
| Average-Remote/Recent | .94*** | .92*** | .86*** | .95*** | .95*** | .83*** | .84*** | .95*** |
| Older Adults | Internal Count | Internal Density | Proportion Score | External Count | Semantic Count | External Density | Semantic Density | Word Count |
| Childhood-Previous Year (Remote-Recent) | .49*** | .19* | .34*** | .34*** | .38*** | .35*** | .23** | .49*** |
| Childhood-Average | .70*** | .54*** | .62*** | .79*** | .76*** | .63*** | .64*** | .78*** |
| Teenage-Average | .74*** | .66*** | .64*** | .82*** | .85*** | .62*** | .66*** | .87*** |
| Early Adult-Average | .83*** | .76*** | .71*** | .85*** | .84*** | .64*** | .69*** | .90*** |
| Middle Adult-Average | .77*** | .71*** | .69*** | .84*** | .86*** | .62*** | .73*** | .89*** |
| Previous Year-Average | .81*** | .74*** | .77*** | .69*** | .72*** | .74*** | .58*** | .78*** |
| Childhood-Remote/Recent | .73*** | .78*** | .77*** | .75*** | .71*** | .80*** | .81*** | .76*** |
| Teenage-Remote/Recent | .56*** | .55*** | .49*** | .61*** | .63*** | .41*** | .44*** | .69*** |
| Early Adult-Remote/Recent | .57*** | .54*** | .51*** | .65*** | .62*** | .40*** | .35*** | .69*** |
| Middle Adult-Remote/Recent | .49*** | .45*** | .48*** | .65*** | .62*** | .39*** | .43*** | .66*** |
| Previous Year-Remote/Recent | .96*** | .77*** | .86*** | .87*** | .92*** | .85*** | .76*** | .94*** |
| Average-Remote/Recent | .87*** | .82*** | .86*** | .89*** | .87*** | .84*** | .77*** | .89*** |

Although detail recollection varied with memory age, as predicted, the observed correlations between timepoints suggest that the process of event recollection, as assessed by the AI, is consistent across event memories, irrespective of memory age. This provides empirical support for the decision to average details over all events.

#### Internal consistency (AI metrics)

We next investigated associations among all AI variables. Among the primary AI metrics, we predicted that internal measures would be positively correlated, and that external and semantic measures would be correlated (although this is largely mandated, as semantic details are included in external scores). We predicted that internal and external counts would be correlated with word count, but that word count would not be associated with ratio or density scores.

When examining correlations among AI variables three primary patterns emerged (see Figure 5 for full results). First, all detail count measures were positively correlated. Second, internal density and ratio scores, which adjust for verbosity, were positively correlated, and both were negatively correlated with the external detail measures (count and density). Third, external measures were positively correlated with each other and word count. All patterns held within age groups (Supplemental Figure 1). Contrary to predictions, internal count and internal density were negatively correlated across participants (pr(348)= -.13, p< .05) and within younger adults (pr(198)= -.21, p< .01), but were uncorrelated within older adults (pr(148)= .02, p= .77).

**Figure 5.**
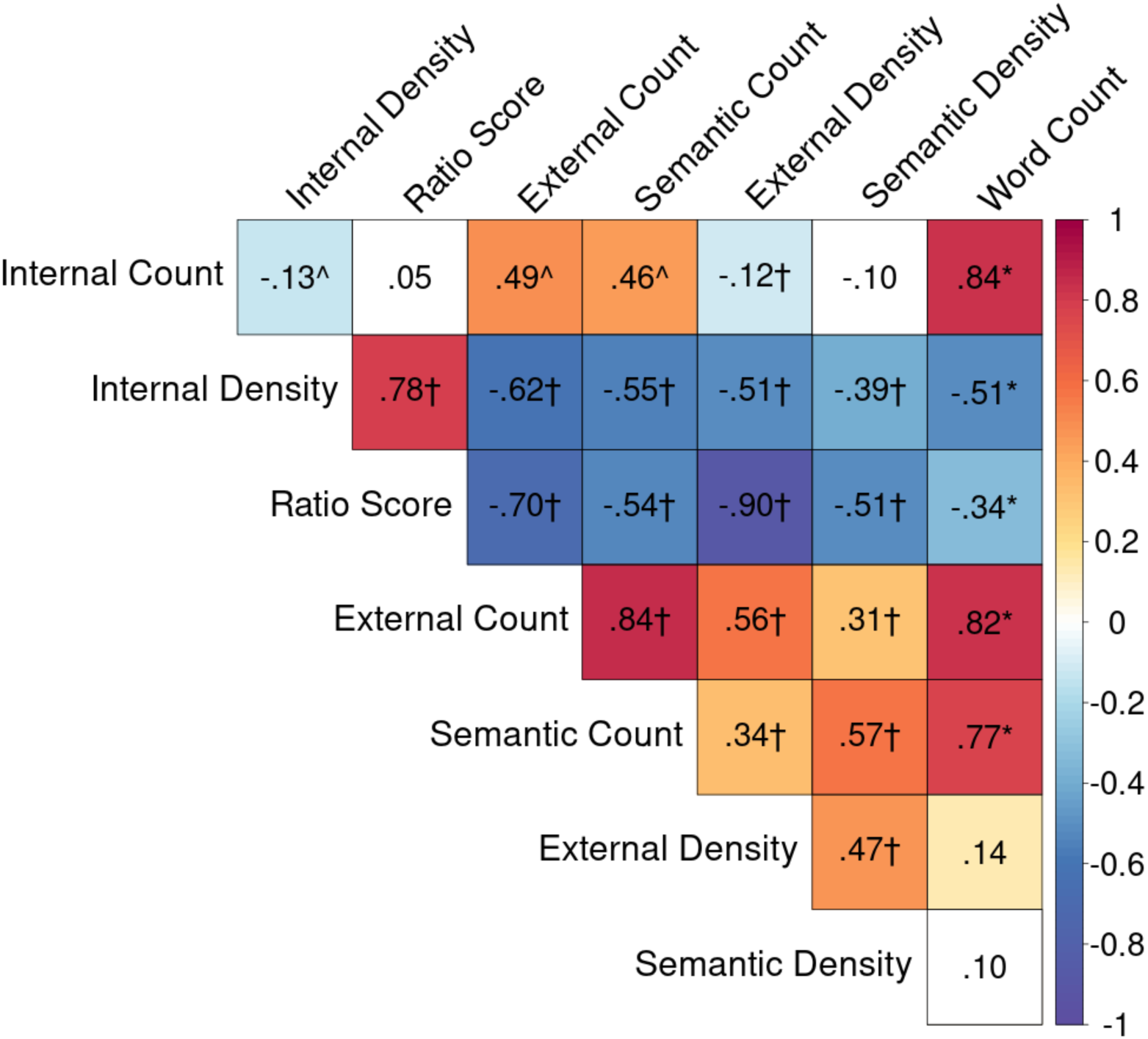
Internal Consistency of the Autobiographical Interview: Measures of Interest. Partial product-moment correlations between the 8 AI measures of interest across all participants controlling for age and gender. The matrix contains r values, which are color-coded by strength of the correlation. Cells with a white background indicate non-significant associations (p > .05, uncorrected). Color cells are significant (p < .05). † significant predicted association, * unpredicted significant association with Bonferroni correction, ^ associations opposite to prediction (p < .05, uncorrected).

Our observation of a positive correlation between internal and external counts is inconsistent with a recent study reporting that these measures were negatively correlated (Devitt et al., 2017). To explore this association further, we conducted a hierarchical linear model analysis (see Methods). The regression weight for internal details was significant (*B* = 0.52, *SE*=0.05, *t* ratio = 11.24, *df* =1130, *p* < .001), indicating that events reported with more internal details were more likely to include more external details. The age group regression weight was also significant (*B* = 2.12, *SE*=0.34, *t* ratio = 6.21, *df*=1160, *p* < .001), indicating that older adults were more likely to provide more external details, as we and others have shown in earlier reports (Levine et al., 2002; Spreng et al., 2018). The interaction term was also significant (*B* = −0.20, *SE*=0.06, *t* ratio =-3.21, *df*=1271, *p* < .01). Internal details were less predictive of external details for older versus younger adults. These analyses provide additional evidence that internal count is positively associated with external (as well as semantic) detail counts. This association remains significant when accounting for age group and within-participant variance.

#### Internal consistency (AI ratings)

The full AI protocol includes self-reported participant ratings and scorer-assigned ratings for level of detail. These self-report and summary rating scores complement the primary focus of the core AI metrics.

We predicted that internal scores would positively correlate with self-report ratings. All ratings were positively correlated with internal count, external count, and total word count (Figure 6). Contrary to predictions, all self-report ratings except rehearsal were negatively correlated with internal density and the ratio score measures. No associations were observed with rehearsal for AI measures that controlled for verbosity (density, ratio). Similar patterns were observed within age groups (Supplemental Figure 2).

**Figure 6.**
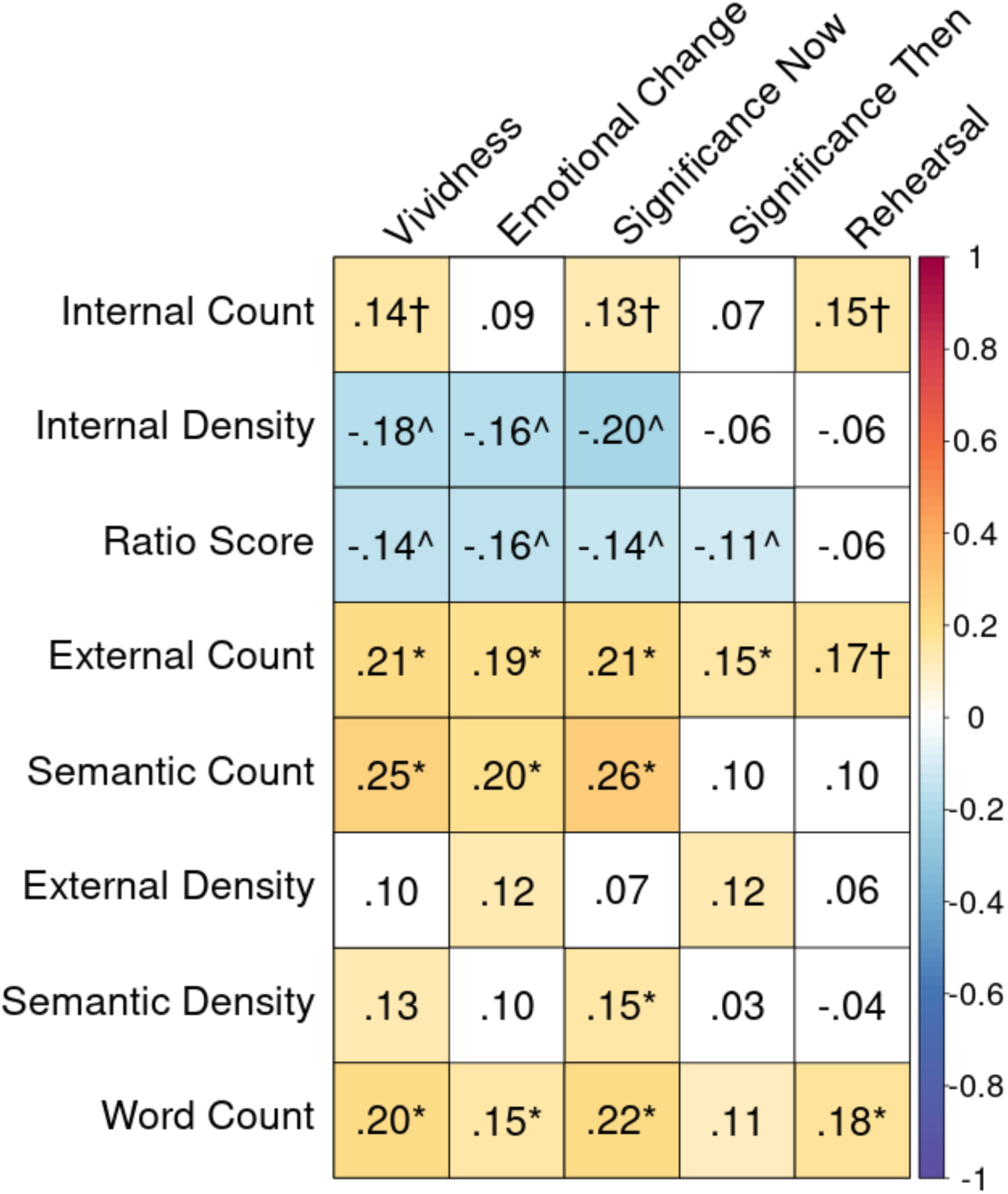
Internal Consistency of the Autobiographical Interview: Participant Self-Report Ratings. Partial spearman correlations between the 8 AI measures of interest and participant self-report ratings across all participants controlling for age and gender. The matrix contains partial ρ values, which are color-coded by strength of the correlation. Cells with a white background indicate non-significant associations (p > .05, uncorrected). Color cells are significant (p < .05). † significant predicted association, * unpredicted significant association with Bonferroni correction, ^ associations opposite to prediction (p < .05, uncorrected

With respect to summary ratings, our predictions follow the scoring guide: internal scores were predicted to associate with ratings that incorporate internal details (place, time, perceptual, emotion/thought, AMI rating, and episodic richness), while external scores were predicted to correlate with ratings that incorporate external details (time integration). Generally, scorer ratings were positively correlated with all counts while associations with density and ratio measures were mixed. Internal measures controlling for verbosity did not show many positive associations with ratings tracking internal detail. Unlike with counts, external density was the only density measure positively correlated with time integration. See Figure 7 for full results. We found similar results within age groups (Supplemental Figure 3) with one notable exception: in younger adults, the AMI rating was negatively correlated with internal density (partial ρ(198)= -.18, p< .01) and ratio score (partial ρ(198)= -.14, p< .05), and episodic richness was negatively correlated with ratio score (partial ρ(198)= -.18, p< .05). In older adults, AMI rating was positively correlated with internal density (partial ρ(148)= .17, p< .05) and ratio score (partial ρ(148)= .42, p< .001), and episodic richness was also positively correlated with the ratio score (partial ρ(148)= .32, p< .001).

**Figure 7.**
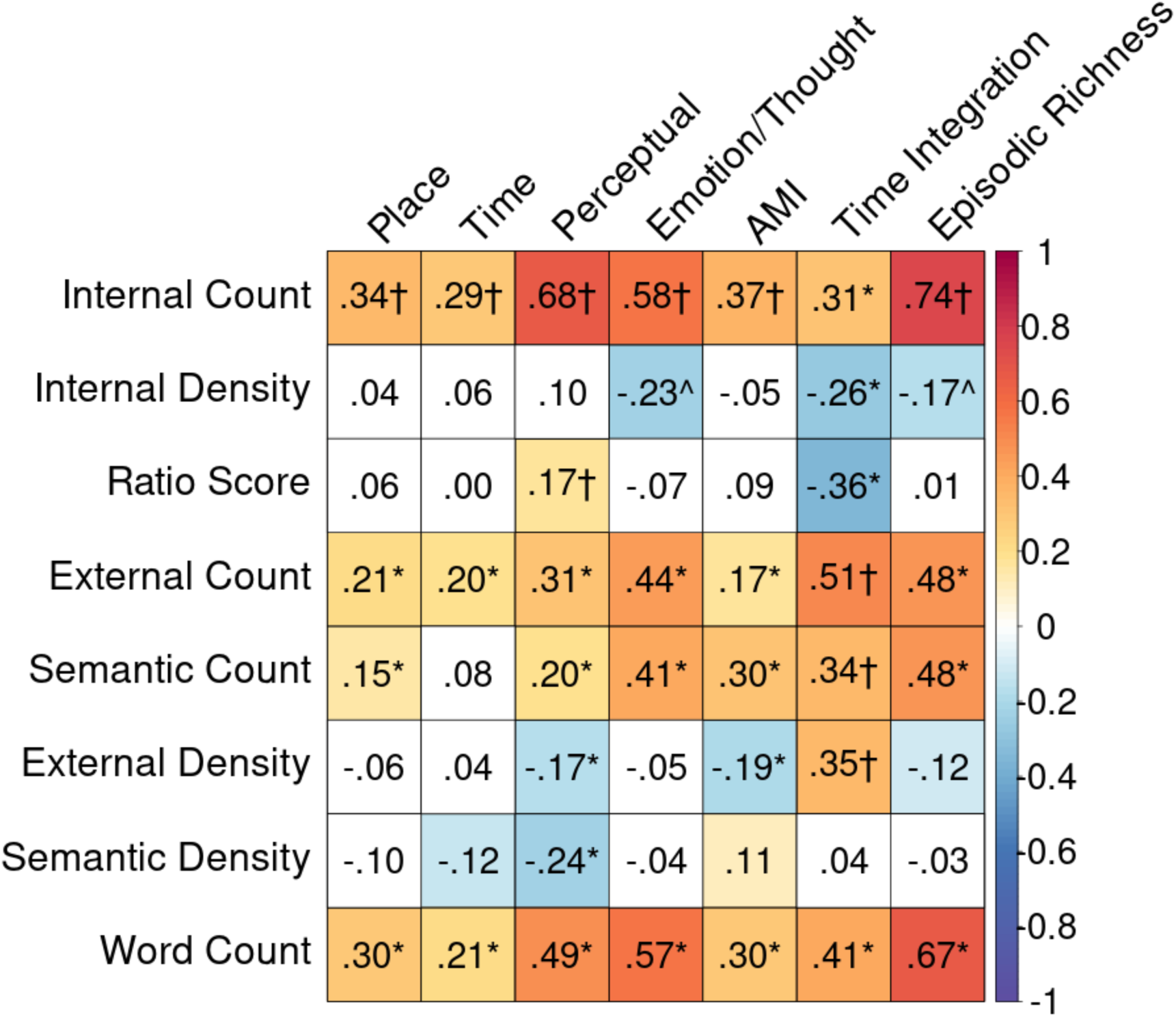
Internal Consistency of the Autobiographical Interview: Scorer Ratings. Partial spearman correlations between the 8 AI measures of interest and average scorer ratings across all participants controlling for age and gender. AMI= episodic specificity rating from the Autobiographical Incident Schedule of the Autobiographical Memory Interview. The matrix contains partial ρ values, which are color- coded by strength of the correlation. Cells with a white background indicate non-significant associations (p > .05, uncorrected). Color cells are significant (p < .05). † significant predicted association, * unpredicted significant association with Bonferroni correction, ^ associations opposite to prediction (p < .05, uncorrected).

### (ii) Demographic Associations with the Autobiographical Interview

We next assessed how AI scores varied by demographics, including age, gender and education (Figure 8). We predicted a robust age effect with higher internal details in younger adults, and higher external details for older adults. Few reports have examined sex and gender differences in AM, and the findings to date have been mixed (e.g., Compere et al., 2016; Fuentes and Desrocher, 2013). Similarly, little work has directly assessed the impact of education on AI performance or AM more broadly. As such, we offer no specific predictions with respect to sex/gender or education differences.

**Figure 8.**
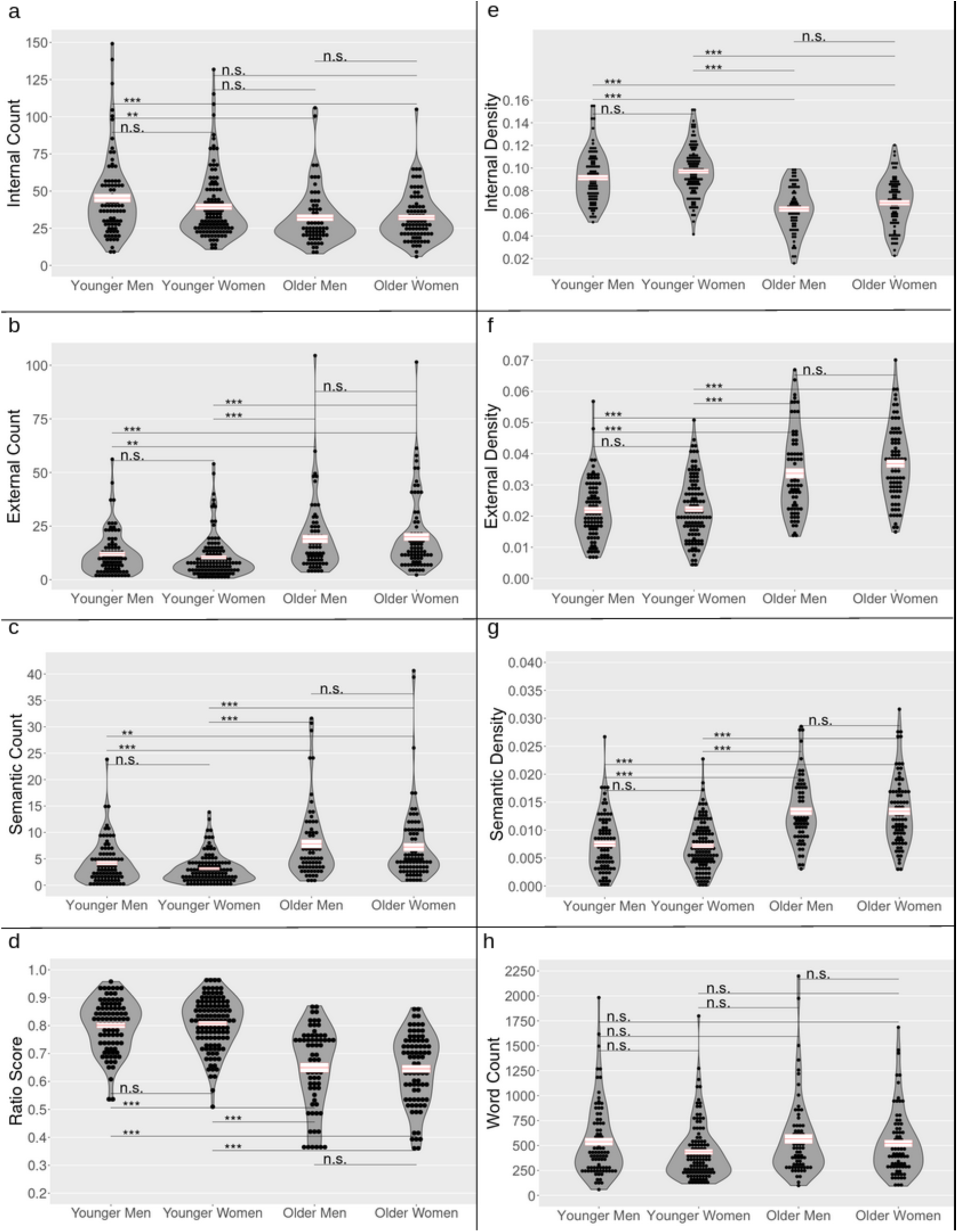
Gender and Age Differences. The a) internal, b) external, and c) semantic counts, d) the ratio score, the density scores for e) internal f) external and g) semantic details, and h) the word count averaged across raters and time periods and separated by gender and age groups. Error bars represent the mean +/- standard error. Significant differences were identified using pairwise t-tests. P values are Bonferroni corrected for tests without a-priori hypotheses; n.s. not significant; *p<.05; **p < .01; ***p<.001.

The predicted main effects of age were supported, with significant age effects observed on all AI measures except for word count. Compared with older adults, younger adults had higher internal count (F(1, 348)=18.11, p<.001, Cohen’s d =0.45), higher internal density (F(1, 348)=144.09, p<.001, Cohen’s d =1.29), and a higher ratio of internal to total details (F(1, 348)=172.23, p<.001, Cohen’s d =1.42). Compared to younger adults, older adults had higher external count (F(1, 348)=35.73, p<.001, Cohen’s d =0.65), higher semantic count (F(1, 348)=48.19, p<.001, Cohen’s d =0.75), higher external density (F(1, 348)=126.60, p<.001, Cohen’s d =1.21), and higher semantic density (F(1, 348)=115.94, p<.001, Cohen’s d =1.16).

We found evidence for a small gender effect (F(1, 348)=7.47, p<.01, Cohen’s d = 0.25), where women had higher internal density than men, although men had a higher overall word count (F(1, 348)=4.81, p<.05, Cohen’s d = 0.23). There were no significant interaction effects (F(1, 348)=.00028-1.61, p>.21).

There is little consensus on the relationship between education and AM with both null (Berna et al., 2012; Murre et al., 2014) and positive associations previously observed (Borrini et al., 1989; Wessel et al., 2014). A modest negative correlation was observed in the current sample between external density and years of education across all individuals (pr(348)= -.12, p< .05), but did not remain significant within either the younger or older adult groups (younger pr(198)= -.13, p=.07; older pr(150)=-.10, p=.22). No other significant associations were observed between AI scores and years of education across all participants or within age groups. Based on the small *r* value with external density and the lack of any other associations, we did not control for education in subsequent analyses (Cohen, 1988; Gignac & Szodorai, 2016).

### (iii) Convergent Validity of the Autobiographical Interview

We next examined the convergent validity of the AI scores with index scores of episodic memory, semantic memory, executive function and the retrieval processes of recognition and familiarity.

We first confirmed that cognitive functions followed expected age patterns (Park et al., 2001). Episodic index scores were significantly higher for young adults (Young: M=0.52, SD=0.46; Old: M=-0.70, SD=0.71; t(241) = 18.38, p < .001, Cohen’s d = 2.10). Semantic index scores were significantly higher for older adults (Young: M=-0.37, SD=0.75; Old: M=0.50, SD=0.71; t(331) = 10.99, p < .001, Cohen’s d =1.18). Executive function index scores were significantly higher for young adults (Young: M=0.35, SD=0.52; Old: M=-0.46, SD=0.51; t(326) = 14.66, p < .001, Cohen’s d =1.57).

Consistent with the established theories of discrete memory systems, and the original framing of the AI as an instrument capable of differentiating episodic and semantic recollection, we predicted that laboratory-based episodic memory measures would positively correlate with internal detail metrics on the AI and that laboratory-based semantic memory measures would positively correlate with external detail metrics. Based on prior work, we anticipated large effect sizes for these effects, with correlation coefficients of approximately .30 (Gignac & Szodorai, 2016; Hemphill, 2003). We also predicted that executive function measures would positively correlate with internal details and negatively correlate with external details, consistent with work highlighting the role of executive function in the search and retrieval of episodic memories (Abellán-Martínez et al., 2019; Conway & Pleydell-Pearce, 2000; Yubero et al., 2011). We anticipated a medium-small effect size for this association, with a correlation coefficient around .15.

Internal AI measures were positively correlated with standard measures of episodic memory. Across the whole sample, the episodic memory index score was positively correlated with internal count (pr(348)= .24, p< .001), internal density (pr(348)= .15, p< .01), and ratio score (pr(348)= .28, p< .001). The episodic memory index score was negatively correlated with external density (pr(348)= -.25, p< .001). The semantic memory index score was not significantly associated with any of the AI measures (pr(341) range= -.06 – .05, *p*’s= .25 – .93). Consistent with predictions, the executive function index score was significantly associated with internal density (pr(348) = .10, p = .05). No other associations were observed with executive function. Correlation between the AI and index scores are shown in Figure 9. Association patterns are also evident in the individual measures that comprise the index scores as visualized in Supplemental Figure 4.

**Figure 9.**
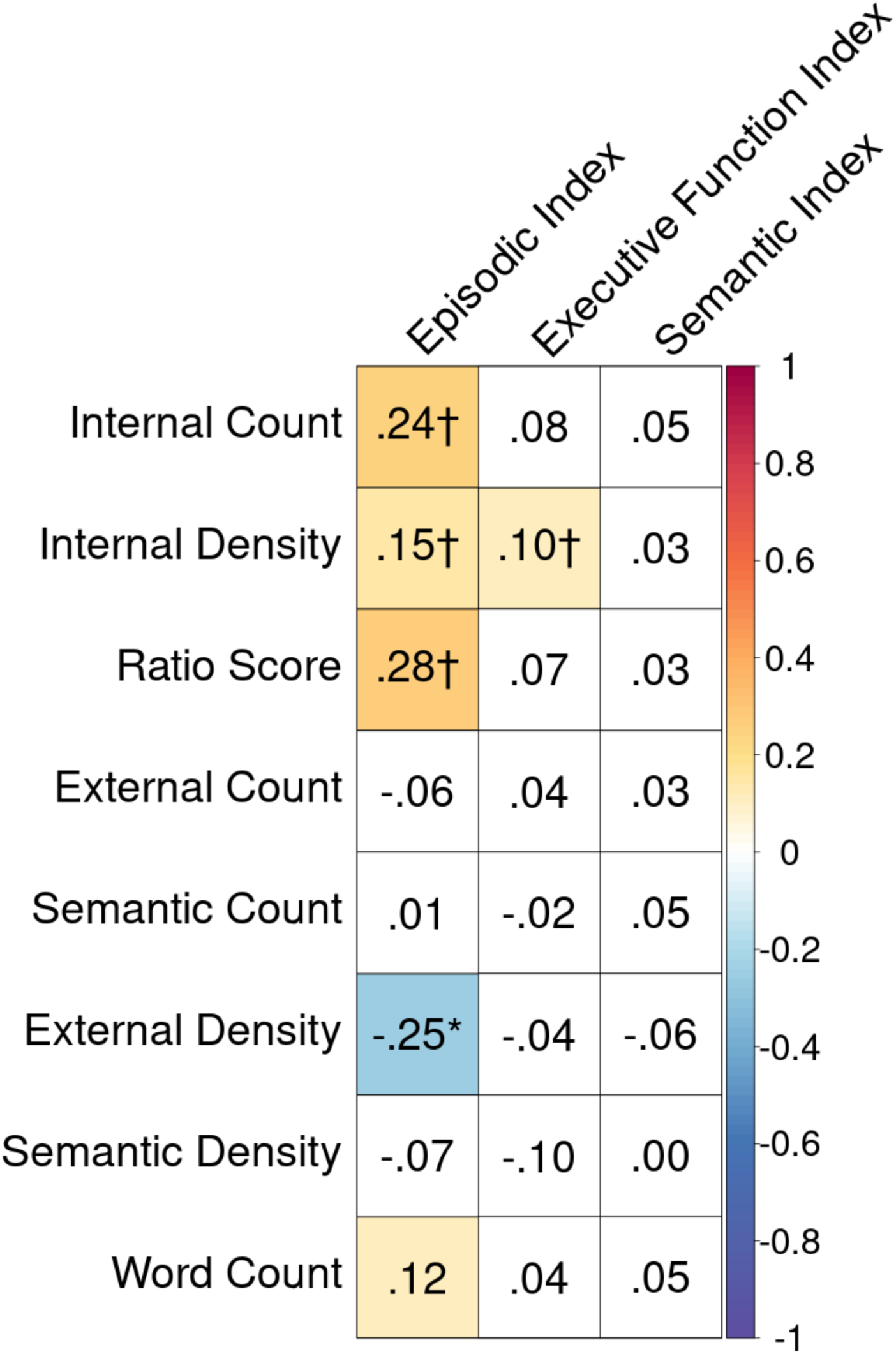
Convergent Validity of the Autobiographical Interview with Episodic Memory, Executive Function, and Semantic Memory Index Scores. Partial product-moment correlations between the 8 AI measures of interest and the 3 index scores across all participants controlling for age and gender. The matrix contains partial r values, which are color-coded by strength of the correlation. Cells with a white background indicate non-significant associations (p > .05, uncorrected). Color cells are significant (p < .05). † significant predicted association, * unpredicted significant association with Bonferroni correction.

We found similar results within age groups, with some exceptions (Supplemental Figure 5). In younger adults, internal density was not associated with either the episodic index (pr(198)= .01, *p*= .90) or the executive function index (pr(198)= .08, *p*= .25) . In older adults, the executive function index was not associated with internal density (pr(148)= .13, *p*= .10) but was negatively associated with semantic density (pr(148)= -.16, *p*<.05). See Supplemental Figure 6 for association patterns between individual measures that comprise the index scores in the younger and older groups.

Drawing from previous work showing that memory specificity is positively associated with recollection and negatively associated with familiarity (Piolino et al., 2006), we predicted that internal detail measures would be positively and specifically correlated with recollection on the R/K task, as they putatively assess episodic re-experiencing. Contrary to our predictions, no associations with recollection were observed for the full sample (Figure 10). Negative associations were observed between familiarity and external count (pr(234)= -.13, *p*< .05), semantic count (pr(234)= -.20, *p*< .01), and semantic density (pr(234)= -.18, *p*< .01). In contrast, the ratio score was positively correlated with familiarity (pr(234) range= .18, *p*< .01). No other associations were significant across the full sample. No significant correlations emerged within the younger adult group. For older adults we observed an additional positive correlation between familiarity and internal density (pr(87)= .27, *p*< .01).

**Figure 10.**
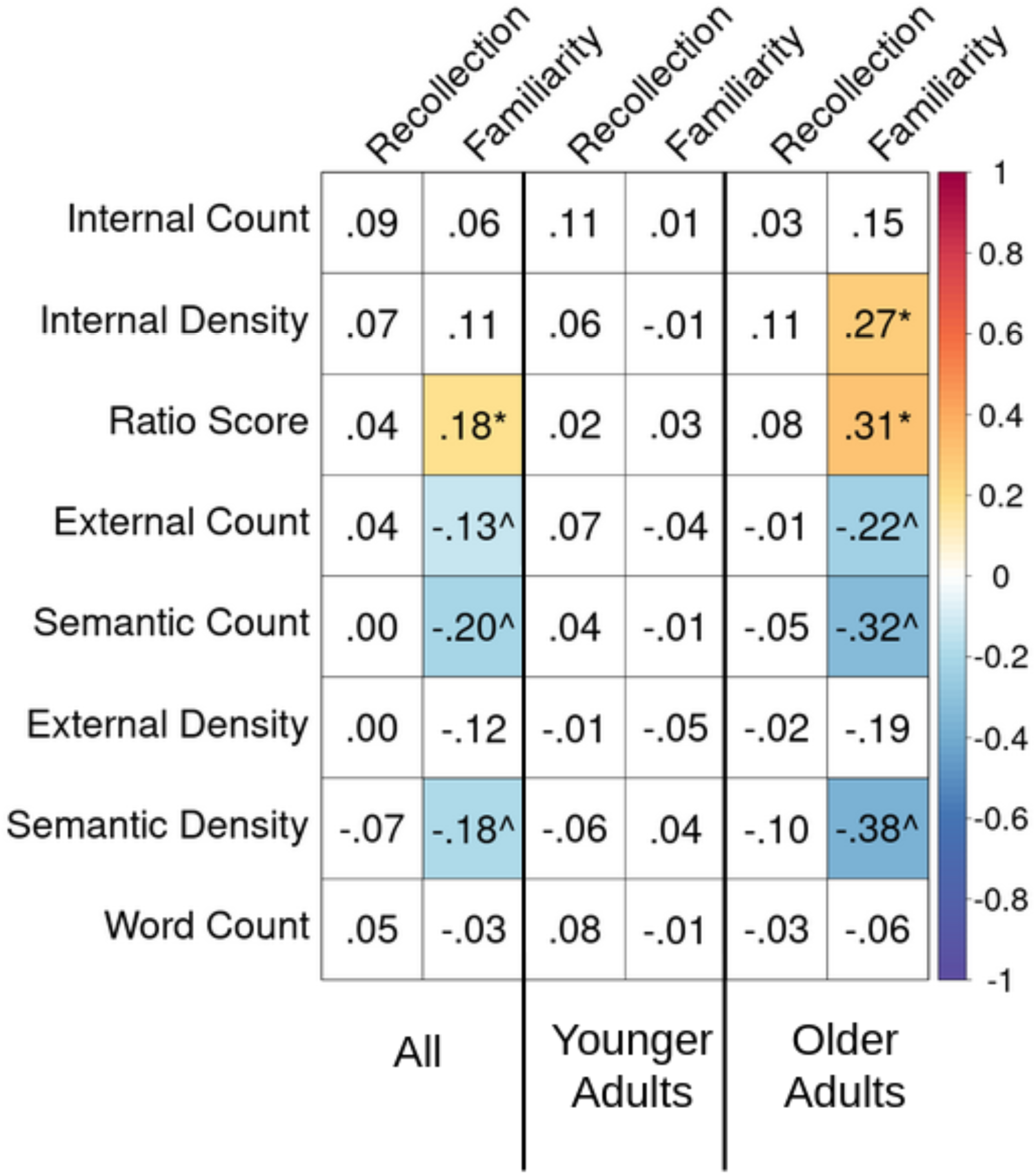
Convergent Validity of the Autobiographical Interview with Measures of Recollection and Familiarity. Partial product-moment correlations between the 8 AI measures of interest and measures of recollection and familiarity from the Remember-Know paradigm across all participants controlling for age and gender, in younger adults controlling for gender, and in older adults controlling for gender. The matrix contains partial r values, which are color-coded by strength of the correlation. Cells with a white background indicate non-significant associations (p > .05, uncorrected). Color cells are significant (p < .05). † significant predicted association, * unpredicted significant association with Bonferroni correction, ^ associations opposite to prediction (p < .05, uncorrected)

### (iv) Associations with Depression, Decision-Making, Social Cognition and Personality

Based on previous reports implicating non-mnemonic factors in AI performance, we examined associations between AI and measures of depression, decision-making (temporal discounting), social cognition, and personality. We anticipated small to typical effect sizes, with correlation coefficients around .10-.20 (Gignac & Szodorai, 2016; Hemphill, 2003) given the indirect but robust links between AM and our non-mnemonic domains of interest.

### Depression

We predicted that internal measures would be negatively correlated with measures of depression, given previous reports of an inverse relationship between low mood and memory performance (Brittlebank et al., 1993; Hitchcock et al., 2014; Kuyken & Dalgleish, 1995; Liu et al., 2013; Williams & Broadbent, 1986; Williams & Scott, 1988; Williams et al., 2007; Wilson & Gregory, 2018). Because our mood measures differed for younger and older adults (BDI, GDS), we analyzed these associations in each group separately.

In younger adults, depression scores were negatively associated with ratio score (pr(115) = -.19, p < .05) and positively associated with external count (pr(115) = .22, p < .05) and external density (pr(115) = .18, p < .05). No significant associations were found between the AI and depression scores in older adults. Correlations between the AI and measures of depression are shown in Figure 11.

**Figure 11.**
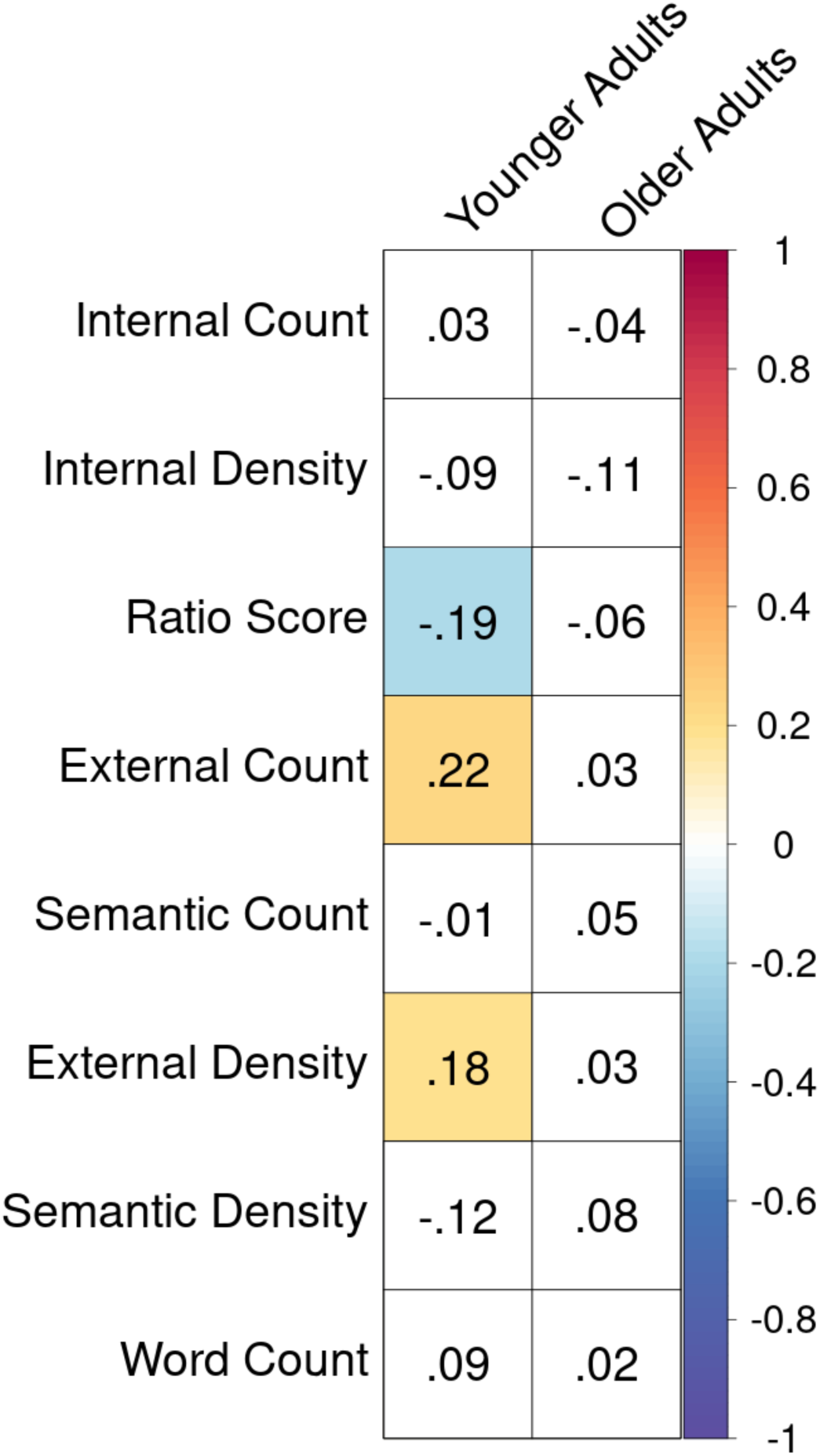
Convergent Validity of the Autobiographical Interview with Measures of Depression. Partial product-moment correlations between the 8 AI measures of interest and the Beck Depression Inventory in younger adults and the Geriatric Depression Scale in older adults across participants controlling for gender. The matrix contains partial r values, which are color-coded by strength of the correlation. Cells with a white background indicate non-significant associations (p > .05, uncorrected). Color cells are significant (p < .05). † significant predicted association, * unpredicted significant association with Bonferroni correction.

### Decision-making (temporal discounting)

We predicted that internal detail measures would be negatively correlated with temporal discounting based on recent work by Lempert and colleagues (2020), which suggested that the propensity to wait for larger rewards is related to episodic AM capacity. We did not observe an association between temporal discounting and AI measures across participants or within age groups (Supplemental Figure 7). Temporal discounting was not associated with the ratio of perceptual/gist-based recollection reported in Lempert et al. (2020).

### Social Cognition

For measures of social cognition, we predicted positive correlations with internal detail measures given previously reported associations between theory of mind, sociality and AM (Buckner & Carroll, 2007; Buckner et al., 2008; Gaesser, 2013; Gaesser & Schacter, 2014; DuPre et al., 2016; Rabin et al., 2010; Spreng et al., 2009; Spreng & Grady 2010; Spreng & Mar, 2012). Consistent with predictions, across all participants, performance on the Reading of the Mind in the Eyes task was positively associated with ratio score (pr(300)= .16, p < .01) and negatively correlated with external density scores (pr(300)= -.21, p < .01; Figure 12). A similar pattern was observed in younger adults (Supplemental Figure 8a). For older adults, associations with a second measure of social cognition emerged. Perspective taking on the Interpersonal Reactivity Index was positively correlated with internal count (pr(132)= .19, p < .05) and semantic count (pr(132)= .24, *p* < .01; Supplemental Figure 8b).

**Figure 12.**
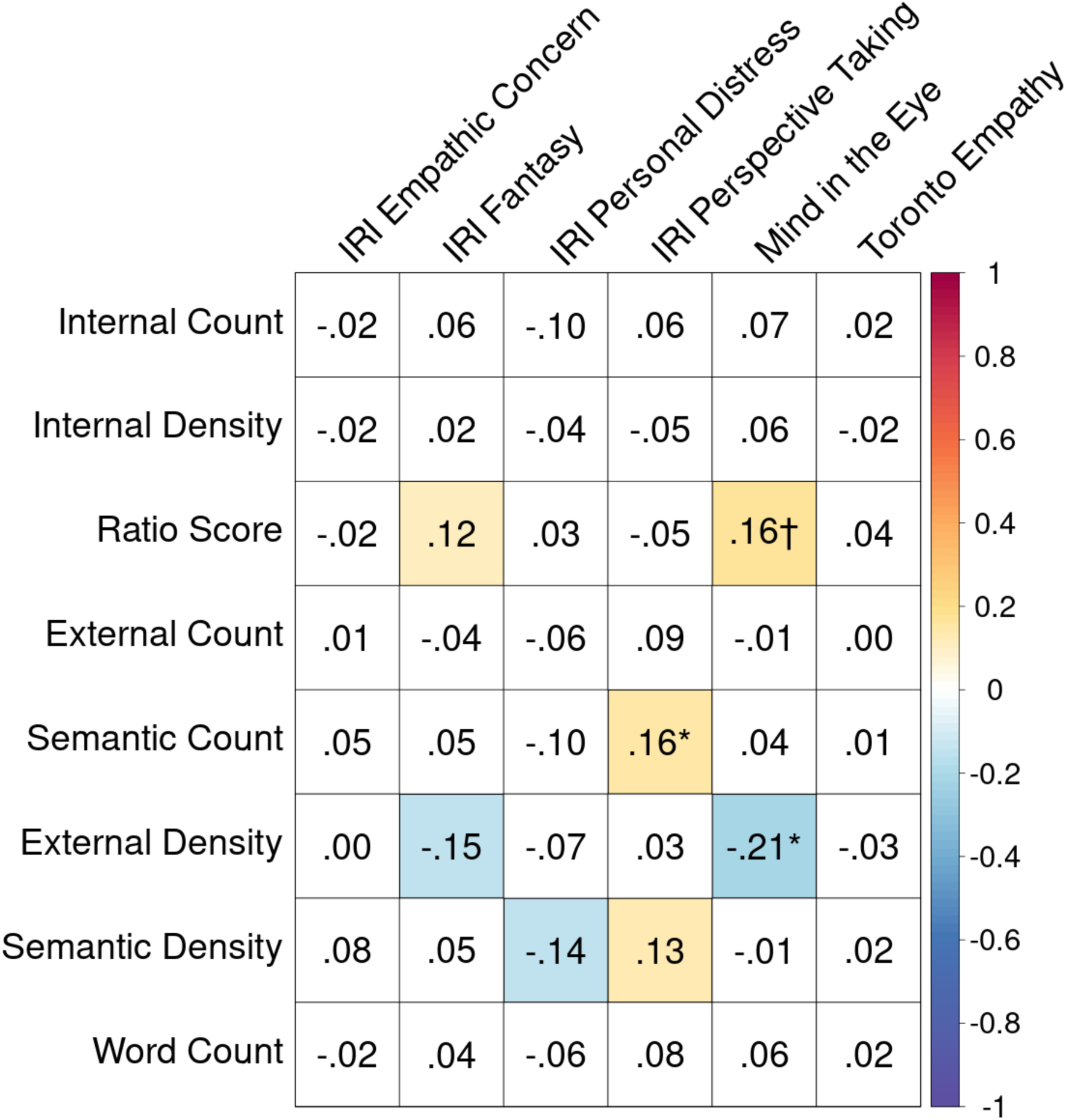
Associations between the Autobiographical Interview and Measures of Social Cognition. Partial product-moment correlations between the 8 AI measures of interest and measures of social cognition across all participants controlling for age and gender. The matrix contains partial r values, which are color-coded by strength of the correlation. The matrix contains r values, which are color-coded by strength of the correlation. Cells with a white background indicate non-significant associations (p > .05, uncorrected). Color cells are significant (p < .05). † significant predicted association, * unpredicted significant association with Bonferroni correction

### Personality

We predicted that internal detail measures would be positively correlated with trait openness/intellect based on previous work linking trait openness with autobiographical coherence, narrative complexity, vividness, sensory experience, and reliving (Adler et al., 2007; McAdams et al., 2004; Rasmussen & Berntsen, 2010; Rubin & Siegler, 2004). No associations survived Bonferroni correction across participants (Figure 13) or in younger adults (Supplemental Figure 9a). In older adults (Supplemental Figure 9b), openness/intellect was positively associated with internal count (pr(133)= .18, p< .05) and ratio score (pr(133)= .17, p< .05).

**Figure 13.**
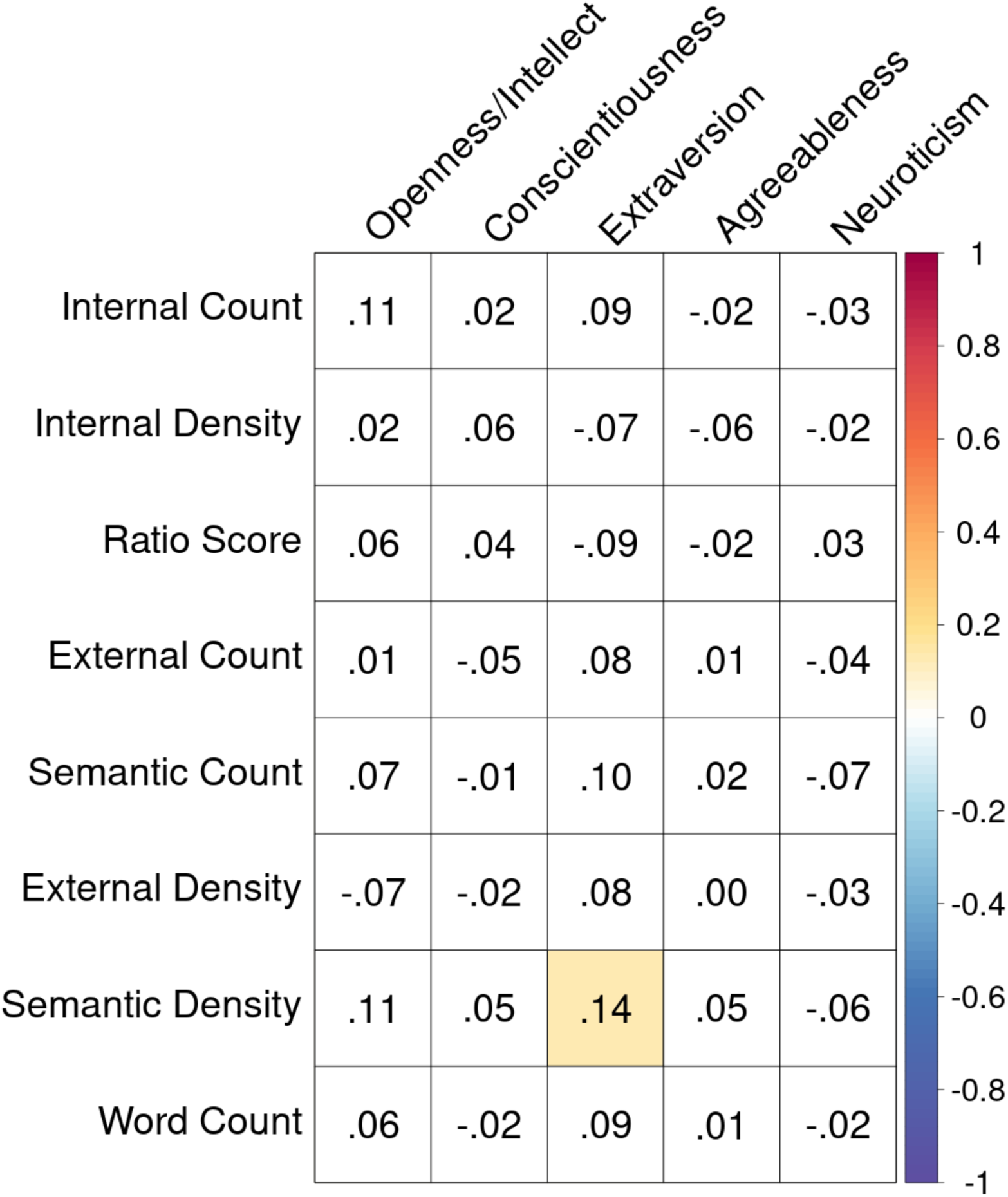
Associations between the Autobiographical Interview and Measures of Personality. Partial product-moment correlations between the 8 AI measures of interest and measures of personality across all participants controlling for age and gender. The matrix contains partial r values. Cells with a white background indicate non-significant associations (p > .05, uncorrected). Color cells are significant (p < .05).

### (v) Factor Analyses

#### Confirmatory Factor Analysis

Based upon the intended design of the AI to dissociate episodic from semantic features of AM, we predicted that a two-factor model of the AI would be supported by the data. However, a two-factor model for internal and external details was not supported in the CFA. Low model fit was observed for both count and density metrics.

The CFA with detail counts showed a low model fit: CFI=.76, TLI= .71, RMSEA= .155 (90% CI = [.143, .167]). Recommended thresholds for good model fit are CFI at or above .95 to surpass performance of a null model (Hooper et al., 2008), TLI at or above .95 (Hooper et al., 2008), and RMSEA at or below .08 (Browne & Cudeck, 1993). This RMSEA value indicates a large difference between the residuals of the sample covariance matrix and the hypothesized covariance model. Observed variables all showed significant positive factor loadings (see Supplemental Figure 10). The two latent variables (internal/external) were also positively correlated in younger adults (r(199)=.78, *p*< .001) and in older adults (r(149)= .37, *p*< .001).

The CFA with density scores also showed a low model fit: CFI= .64, TLI= .56, RMSEA= .130 (90% CI = [.118, .142]). See Supplemental Figure 11 for factor loadings. The two latent variables (internal/external) were negatively correlated in younger (r(199)= -.29, *p*< .01) and in older adults (r(149)= -.50, *p*< .001), broadly consistent with a dissociation between internal and external features of AM. A Vuong closeness test suggested that the CFA model with density scores was closer to the true data generating process than the CFA with counts (z = 154.18, *p* < .001).

Given the potential effect of including non-mnemonic content in the external detail category, we ran one CFA without repetition and other details in the model. Neither the count (CFI=.85, TLI= .80, RMSEA= .133 (90% CI = [.118, .147]) nor the density models (CFI=.74, TLI= .66, RMSEA= .126 (90% CI = [.111, .140]) demonstrated acceptable model fit. Associated factor loadings are visualized in Supplemental Figures 12 and 13.

#### Exploratory Factor Analysis

EFA analyses were conducted to identify potential alternatives to the two-factor AI model. We found variation in the optimal number of factors recommended by the Parallel Analysis procedure within different demographic subsets: supported models ranged from two to four factors (Figure 14 and Supplemental Figures 14 and 15).

**Figure 14.**
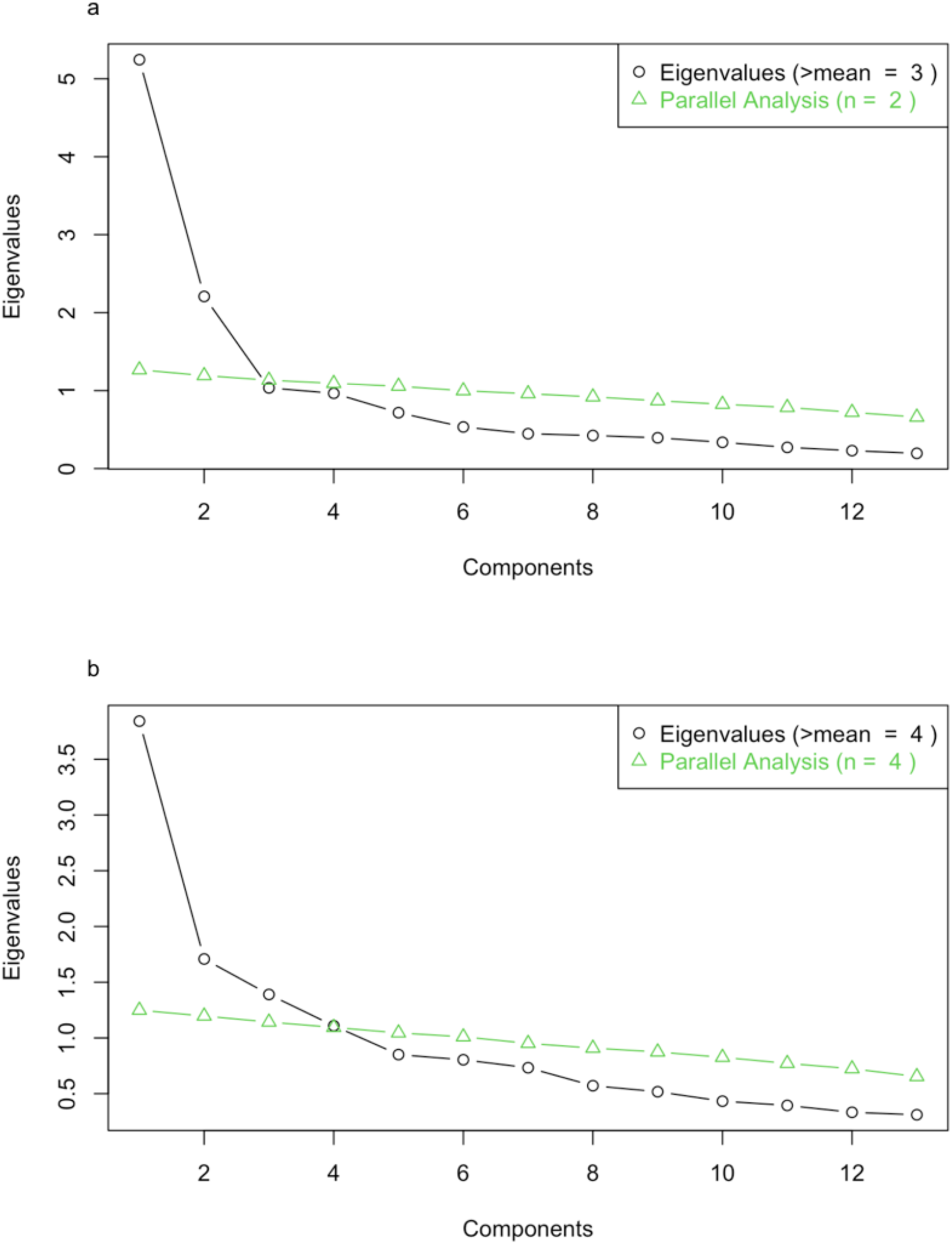
Exploratory Factor Analysis of the Autobiographical Interview Across All Participants. Resulting eigenvalues and parallel analysis results for an optimal number of factors from an exploratory factor analysis of a) detail counts and b) density scores.

Across all participants, parallel analysis identified a two-factor model for count measures that accounted for 50% of the overall variance. Eigenvalues for each factor, or component, are displayed in Figure 14a. These factors loaded discretely onto internal and external details, with the exception of semantic, repetition, and other details which loaded onto both factors. Factor loadings are shown in Table 4. In contrast, a four-factor model emerged with density scores, accounting for 46% of the overall variance (Figure 14b). Detail categories were roughly separated into internal, external, semantic, and semantic/repetition/other categories (Table 5).

**Table 4.** Factor Loadings for Exploratory Factor Analysis of Detail Counts Across All Participants. Factor loadings from exploratory factor analysis with varimax rotation for each of two factors onto detail sub-categories. Factor columns are sorted by variance explained (most to least). Loadings greater than .3 are bolded.

| Detail type | Factor loading |  |
| --- | --- | --- |
|  | 1 | 2 |
| External Event Details | **.85** | .13 |
| External Perceptual Details | **.76** | .16 |
| External Place Details | **.72** | .01 |
| External Emotional Details | **.68** | .12 |
| External Time Details | **.62** | .17 |
| External Semantic Details | **.58** | **.40** |
| External Other Details | **.53** | **.51** |
| External Repetitions | **.40** | **.54** |
| Internal Perceptual Details | .02 | **.82** |
| Internal Event Details | .12 | **.79** |
| Internal Emotional Details | .22 | **.63** |
| Internal Place Details | .10 | **.58** |
| Internal Time Details | .08 | **.42** |

**Table 5.** Factor Loadings for Exploratory Factor Analysis of Density Scores Across All Participants. Factor loadings from exploratory factor analysis with varimax rotation for each of four factors onto detail sub-categories. Factor columns are sorted by variance explained (most to least). Loadings greater than .3 or less than -.3 are bolded.

| Detail type | Factor loading |  |  |  |
| --- | --- | --- | --- | --- |
|  | 1 | 2 | 3 | 4 |
| External Event Details | **.83** | -.12 | -.12 | .19 |
| External Perceptual Details | **.70** | **.30** | -.10 | -.03 |
| External Emotional Details | **.59** | -.06 | -.27 | .09 |
| External Place Details | **.58** | .21 | .03 | .25 |
| External Time Details | **.45** | .23 | .03 | .24 |
| External Other Details | .09 | **.65** | -.06 | .11 |
| External Repetitions | .03 | **.49** | -.09 | .07 |
| Internal Place Details | -.20 | -.08 | **.74** | -.04 |
| Internal Time Details | -.05 | -.24 | **.62** | -.09 |
| Internal Event Details | -.24 | **-.55** | **.36** | **-.34** |
| External Semantic Details | .02 | **.30** | -.20 | **.65** |
| Internal Emotional Details | -.12 | -.10 | -.17 | -.28 |
| Internal Perceptual Details | -.26 | .00 | .14 | **-.53** |

In younger adults only, parallel analysis identified a three-factor model for count measures, accounting for 52% of the overall variance. Loadings did not appear to separate along any clear categorical lines (Supplemental Table 1). With density scores, a four-factor model emerged, accounting for 41% of the overall variance (Supplemental Figure 14). This model was roughly separated into factors of internal details (except emotion/thought), external details, semantic details, and semantic/repetition/other details (Supplemental Table 2).

In older adults only, parallel analysis identified a two-factor model for count measures, accounting for 56% of the overall variance. This model divided internal from external details with the exception of semantic, repetition, and other categories, which shared variance across the two factors (Supplemental Table 3). For density scores, a three-factor model accounted for 43% of the overall variance (Supplemental Figure 15). This model loaded factors according to internal details, most external details, and repetition/other details (Supplemental Table 4).

## Discussion

The AI is a performance-based measure of AM that has emerged as the gold standard for quantifying and characterizing personal recollective experiences. Despite the prominence of the AI, there has yet to be a comprehensive evaluation of its psychometric properties. Here we report the findings of a psychometric assessment of the AI in a well-powered sample of healthy younger and older adults. AI performance is reported using both the original approach (detail counts and ratio scores; Levine et al., 2002) and novel metrics wherein detail counts are scaled by verbosity (density scores). Our first aim was to determine whether the AI was a reliable and valid measure of AM. Inter-rater reliability and internal consistency suggested high reliability, but this depended on the metric used. Density scores were more stable across recalled events. Reliability was further confirmed with robust age effects across detail counts, ratio scores, and density scores. Other demographic factors had more modest effects and appear to be less replicable across samples. Convergent validity analyses provided strong evidence that the AI is a valid measure of episodic, but not semantic AM. Performance on standard episodic memory tasks was correlated with all internal detail metrics. No associations were observed with standard semantic memory measures. Density scores also correlated with executive function measures, suggesting added sensitivity of verbosity-controlled metrics to AM processes. Given the rich literature on non-mnemonic influences on AI performance, our second aim was to replicate previously reported associations between the AI and individual difference measures of psychological and cognitive function. Associations between depression and the AI were observed in the younger adults. Few other predicted associations with these non-mnemonic factors were observed. An important theoretical assumption underlying the design of the AI is that separate scoring of internal and external details disambiguates episodic from non-episodic event content. Our third aim was to empirically test this assumption. The two-factor model was not strongly supported by our data, but model fit was improved with density scores. EFA provided alternative factor structures for exploration in future work. As each of the key analyses in our assessment was performed on original and novel AI metrics, our final aim was to evaluate the merits of controlling for verbosity. Overall, we find that controlling for verbal output reveals a more stable pattern of AM performance, increased convergent validity with standard laboratory tasks of cognitive functioning, and improved correspondence to the proposed two-factor model of AM. We conclude by discussing the importance of considering verbosity when interpreting performance on the AI.

### The Autobiographical Interview is highly reliable

#### Inter-rater reliability

A significant challenge for quantifying AM is the objective evaluation of another’s memory. Determining what comprises a recollected event and the identification of information units within an event narrative is subjective. However, our findings demonstrate that the AI’s rigorous administration and scoring protocol are highly effective at reliably parsing recollective experiences into internal and external details. This is critically necessary for characterizing and quantifying the episodicity of a recollected experience (Levine et al., 2002). Inter-rater reliability on the AI scoring was high, far exceeding established benchmarks in the field (e.g., .75-.80; Cicchetti, 1994; Orwin, 1994). The inter-rater reliability we observe is comparable to previously reported intraclass correlation values in smaller samples, which range from .88-.96 for internal details and .79-.96 for external details (Addis et al., 2008; Addis et al., 2010; Cole et al., 2012; Devitt & Schacter, 2020; Gaesser et al., 2011; Levine et al., 2002; Robin & Moscovitch, 2017; Terrett et al., 2016). While not the focus of the current report, inter-rater reliability was similarly high for internal detail sub-categories as well as for semantic details. These findings suggest that core features of autobiographical re-experiencing can be reliably quantified using the AI.

#### Internal consistency (across timepoints)

We observed high internal consistency across events recalled from different time periods. While AM clearly varies as a function of time (e.g. Conway & Pleydell-Pearce, 2000; Crovitz & Schiffman, 1974; Schroots et al., 2004), our findings suggest that stable individual differences exist in AM capacity that are observable across all time periods. As such, averaging measures across events may be an effective strategy for individual differences research involving the AI. Many studies with the AI compute averages or sums of detail counts across memories (e.g. Fuentes & Desrocher, 2013; Levine et al, 2002; Murphy et al., 2008; Spreng et al., 2018). The present work offers justification for the use of a single metric for healthy younger and older adults. Notably, a robust recency effect was found for count metrics, whereas, verbosity corrected measures did not increase with event recency (although some differences were observed in older adults). Verbosity corrected measures (e.g., density and ratio) may be more appropriate for examining trait-level AM, as we discuss in greater detail below. Robust positive associations between individual events and with composite scores provides strong evidence that each event is measuring a similar latent construct. This finding also suggests that one event may offer sufficient estimation of AM capacity. If validated, this would broaden the feasibility of implementing the AI across a larger range of studies where time and/or sample sizes have precluded its usage.

#### Internal consistency (AI metrics)

A core motivation for the development of the AI was to move beyond one-factor self-report scales of AM capacity such as the Autobiographical Memory Interview (Kopelman et al., 1989, 1990). In contrast, the AI was developed to quantify and dissociate episodic memory (internal details) from other components contributing to AM (external details), including semantic memory, from event recollections. Quantifying this distinction is fundamentally necessary to isolate and thus accurately characterize the episodicity of autobiographical event recall. Here we investigated whether the relationship between internal and external details on the AI dissociates episodic and semantic memory, and reflects a precise characterization of autobiographical re-experiencing. We queried all 8 AI metrics (detail counts, density scores, and ratio scores) to determine whether the relationship was contingent on the dependent variable used. Internal and external counts were positively correlated and were, unsurprisingly, also highly correlated with the number of words spoken. These associations were confirmed by our hierarchical linear modeling analysis. Density and ratio scores revealed a different pattern of associations. Internal density was positively correlated with ratio score, and both were negatively correlated with all external measures, as would be predicted from a two-factor account (internal/external) of AM. We suggest that total words spoken may underlie the association between internal and external detail counts. However, our finding of a positive association between internal and external count is inconsistent with previous works reporting no association (Addis et al., 2008; Addis et al., 2009; Addis et al., 2010; Cole et al., 2012; Gaesser et al., 2011) or a negative association (Devitt et al., 2017). We speculate that this difference may arise from underpowered samples in previous work or variation in AI administration (e.g., time limits for participant responses). The vast majority of AI studies to date has relied on internal count measures (e.g., Palombo et al., 2018; Barry et al., 2020; Armson et al., 2021 for recent examples), but our findings provide evidence that density and ratio scores may better estimate autobiographical episodicity.

The modest negative correlation between internal density and internal count observed here suggests that they measure different constructs and should not be used interchangeably. As mentioned above, it is possible that density scores reflect a more stable, trait-like measure of AM while count scores, sensitive to recency effects, may represent more transient AM abilities. As discussed further below, both measures positively correlate with laboratory episodic memory, which suggests that they both capture some element of episodic AM. We posit that participant verbosity plays a key role in this difference but the link between verbosity itself and phenomenological re-experiencing is unclear. Further research expanding upon the latent structure of these metrics will inform future AI research.

#### Internal consistency (AI ratings)

Full scoring of the AI includes summary ratings of the AI narratives, although these are often omitted from published reports. Scorer ratings include an appraisal of the level of internal place, time, perceptual, and emotion/thought detail provided. Additional summary ratings include the AMI episodic specificity rating, time integration, which conveys the amount of global context provided for the event, and episodic richness, which reflects the impression of participant re-experiencing conveyed by the total amount of internal detail provided.

The AI scoring manual directs scorers to base ratings on internal counts, except for the time integration rating. For this reason, we predicted positive correlations would be observed between these summary ratings and internal count. Consistent with the initial validation of the AI (Levine et al., 2002), internal count was positively associated with the AMI episodic specificity rating. However, most summary ratings were also positively associated with external and semantic count as well as words spoken, with mixed associations observed for verbosity-corrected measures. Time integration was correlated with external density, supporting the use of external density as a measure of non-episodic contextual details. Across the full sample, associations between scorer ratings and AI metrics were non-specific, failing to distinguish between episodic (internal) and non-episodic (external) content. As such, we caution against using scorer ratings in place of recollection-derived internal and external measures.

Participants also rated the quality of their recollections for vividness, emotional change, significance (now and then), and rehearsal. We found that these self-report AI measures were positively associated with all count measures (internal, external, semantic, and word count), as well as semantic density. However, we interpret these associations cautiously as these may reflect non-specific associations with overall verbal output; participants generally speak more about events that they deem to be vivid, emotional, significant, and rehearsed.

In contrast, internal density and ratio scores were negatively correlated with self-reported vividness, emotional change and significance now, mirroring overall positive correlations with external measures. The robust associations between self-report and external details may be evidence for a semantic scaffolding, conceptual integration, or narrative style: higher levels of vividness, emotional change, significance, and rehearsal could be associated with a deeper or more readily available integration into a life-story and self-concept used to access a specific event (Blagov & Singer, 2004; Wood & Conway, 2006). Indeed, these associations were stronger in older adults alone, and were present only with counts in younger adults. In a similar vein, Alzheimer’s disease patients, who provide fewer episodic details during narrative retelling, rate their memories as more significant and emotional (El Haj et al., 2016). Participants may thus provide more semantic, contextual, and metacognitive information to reflect how a particular event fits within their personal semantic landscape (Bluck & Habermas, 2000; Prebble et al., 2013; Addis & Tippett, 2008; Thomsen, 2009). Events that are rated more highly on these self-report measures may be linked to more life themes and knowledge structures that are then shared during the interview. It has been proposed that memories become more semanticized with more rehearsal (Skowronski & Walker, 2004). Consistent with prior work (Fuentes & Desrocher, 2013), external detail count was associated with self-reported rehearsal. Of note, ratio and density scores were unrelated to AM rehearsal. This suggests that ratio and density scores may be more robust to the effects of cuing accessible events versus events that were previously encoded but not previously retrieved (e.g. Svoboda & Levine, 2009).

Substantial debate remains regarding the role of self-reported memory vividness, and its relationship to performance-based AM. As an inherently phenomenological experience, researchers do not have direct access to the recollections of others. Participant reports of vividness have been related to medial temporal lobe activity and connectivity during recollection (Addis et al., 2004; Addis et al., 2011; Furman et al. 2012, Sheldon & Levine 2013; Thakral et al., 2020). However, here we found that vividness was not specifically related to internal count, showing a similar association with external count. In contrast, ratio and density scores showed no relationship with vividness. These observations, and those of others (Clark & Maguire, 2020; Setton et al., 2021), indicate that the AI does not capture many of the multi-faceted phenomenological features of autobiographical recollection (see also Zaman & Russell, 2022).

Further work is needed to more deeply assess the reliability of the AI. Within the same testing session, reliability is high for events sampled from different points in time. Less is known about how test-retest reliability varies over more extended durations. To our knowledge, only a few studies have examined the persistence of AM recollection over time. One study with 12 middle aged adults found that repeated retrieval, more than the passage of time, increased the total of internal and external count (Nadel et al., 2007). The same group also reported that passage of one year led to increased detail recollection (Campbell et al., 2011). A more recent study in 16 younger adults found that internal count was nearly identical across two visits separated by 8 months, with only emotion/thought details receding over time (Barry et al., 2020). Replication in larger, more developmentally diverse samples is needed. Importantly, it is currently unknown how AM recollection is affected by cueing, which may vary as a function of highly novel unrehearsed events versus more accessible and general events (for discussion, Armson et al., 2021). Additionally, it has been observed that internal details can be selectively augmented with a specificity induction (Madore et al., 2014). It is unknown if there is variable susceptibility to specificity induction across participants, and induction’s relationship to stable trait estimation of AM.

### The Autobiographical Interview is impacted by age and gender

Consistent with the initial validation and many subsequent studies, we replicated established age effects in AM (e.g. Addis et al., 2008; Addis et al., 2010; Piolino et al., 2002; Piolino et al., 2006; St. Jacques & Levine, 2007): younger adults provided more internal details and older adults provided more external and semantic details. This typifies a more general trend in cognitive aging wherein fluid and cognitive control abilities decline while crystallized cognition is maintained or enhanced (Craik & Bialystok, 2006; Park et al., 2001; Park & Reuter-Lorenz, 2009; Spreng & Turner, 2019). However, non-mnemonic factors may also explain these age-related differences in performance on the AI. Differences in narrative style and discourse goals have been observed between younger and older adults (Adams et al., 1997; Carstensen et al., 1999; James et al., 1998). Similar patterns of age differences in reporting episodic and semantic details have been observed on a perceptual task, in the absence of memory demands (Gaesser et al., 2011). To begin to disentangle memory from narrative style, we previously introduced AI measures corrected for verbosity (Spreng et al., 2018). Here we find that the observed age differences in the AI extended to, and were even larger for, verbosity-corrected measures (ratio score, internal density, external density, and semantic density) indicating that AI age differences are not due to differences in number of words spoken. As we discuss below, this also suggests that ratio score and density measures may be more sensitive to age differences than detail count measures.

Previous findings of gender effects on the AI are mixed. Some studies have reported gender differences (Fuentes and Desrocher, 2013), while other studies have reported no sex differences (Compère et al., 2016). Here we observed higher internal density scores for women, although men were more verbose overall when describing their memories. Previous research on gender differences in AM narrative length has been inconsistent (Grysman & Hudson, 2013). Gender differences have been reported (Niedźwieńska, 2003), although these differences may depend on the degree of retrieval support (Fuentes & Desrocher, 2013). Consistent with our findings, longer memory narratives have been reported in men (Grysman & Denney, 2017), as have more off-topic statements (Baron & Bluck, 2009). However, adolescent and young adult women have also been observed to provide longer narratives (Bohn & Berntsen, 2008; Bohanek & Fivush, 2010; Fivush & Schwarzmueller, 1998; Grysman, 2017; Pillemer et al., 2003). In line with our findings of greater internal density in women, there is evidence that women elaborate further and provide more specific details during AM recollection (Bloise & Johnson, 2007; Grysman, 2017; Grysman et al., 2016; Grysman et al., 2020; Hayne & MacDonald, 2003; Seidlitz & Diener, 1998; Fivush et al., 2012).

### The Autobiographical Interview demonstrates good convergent validity

A primary interest of the present work was to evaluate the convergent validity of the AI by examining its relationship with other theoretically related cognitive constructs. Central to this validation, we predicted a positive association between internal AI metrics and episodic memory. Consistent with our predictions, we found that episodic memory was positively associated with all three internal measures across participants: internal count, ratio score, and internal density. The association between episodic memory performance in the laboratory and AI scores was smaller than anticipated based on prior work, but likely reflects a truer effect size given our larger sample. This convergent validity provides strong support for the AI internal measures, as they track with laboratory-based episodic memory performance (even though internal count is not positively associated with ratio and internal density scores). Notably, internal density was robustly associated with the episodic index score in older adults, but not in younger adults who tended to perform very well with less variability across subjects (although internal density was positively associated with the specific VPA recognition task in both samples as shown in Supplemental Figure 6). We also found that episodic memory was negatively correlated with external density across the full sample and separately within younger and older adults. This finding supports the conceptualization of external density as a measure of non-episodic information produced in the absence of episodic information.

Executive function is necessary for strategic memory retrieval (Conway & Pleydell-Pearce, 2000; Moscovitch, 1992), staying on task (Garon et al., 2008; Fisher & Kloos, 2016), and inhibiting irrelevant information (Ploner et al., 2001; Gazzaley et al., 2005). We predicted that executive function would be positively associated with internal measures, and negatively associated with external measures. We observed a small yet significant association between internal density and executive function, but no other significant associations emerged. This association was largely driven by reading span, which was positively correlated with both ratio and internal density measures. More so than associations with episodic memory, effect sizes relating the more distant constructs of executive function and AM were very small in our sample of healthy adults. It is possible that the association with executive function was tempered by not imposing a time constraint on participants’ recollections and/or the availability of memory retrieval support. Additional work that focuses on the specific elements of executive function involved in AM may strengthen these associations (Gruler et al., 2019).

AM comprises both episodic and semantic features, and the AI aims to capture both with internal and external metrics. Thus, we assessed the relation between AI measures and semantic memory estimated by vocabulary size, a standard measure of crystalized cognition. Although we predicted that external AI measures would be positively associated with our semantic memory index, no associations were found. We contend that this is not a weakness of the AI. Rather, we urge caution when interpreting the external detail categories, which span a wide variety of representational knowledge and personal semantics, both of which are prominent during AM recollection (Renoult et al., 2012; Renoult et al., 2016). Novel scoring procedures for the AI have been introduced to better characterize semantic memory (e.g., Strikwerda-Brown et al., 2019; Renoult et al., 2020), which may demonstrate clearer links between particular semantic sub-categories and the multi-faceted construct of semantic knowledge. Furthermore, the AI administration procedure requires participants to generate specific past events with a focus on recalling internal details. The production of non-specific event information is discouraged. In line with this idea, recent work has found that when participants are asked to describe opinions about everyday events, there is a rise in the number of external details and a reduction in the number of internal details, suggesting that AI details shift as a function of task demands (Strikwerda-Brown, Lévesque, Brambati & Sheldon, 2021). In the absence of any specific cues, individuals may default to schema-based interpretation and reconstruction of events: general impressions of events tend to be remembered first and serve as the basis for additional remembering (Bartlett, 1932). Better measurement of semantic AM may therefore require instructions with less of an episodic focus and/or more of an impressions-based focus (e.g., Madore et al., 2014).

In a final test of convergent validity, we examined relationships between the AI and the R/K task (Tulving, 1972, 1985; Yonelinas et al.,1995), a laboratory measure that aims to dissociate two types of recognition memory processes: recollection and familiarity. Recollection is thought to reflect more qualitative, episodic, high confidence remembering whereas familiarity is thought to reflect the subjective experience of ‘knowing’ in the absence of specific remembering (see Yonelinas, 2002 for review of dual-process models). In this assessment of convergent validity, the AI did not perform as predicted. No associations were observed for recollection, which were hypothesized to relate to internal measures. Contrary to a prior report (Piolino et al., 2006), familiarity was negatively associated with external measures. Participant insight into phenomenological aspects of recollection is a challenge to assess and may have been poorly characterized in this sample. Previous work has reported high rates of “Remember” responses for items that had not previously been seen (Odegard & Lampinen, 2004). Additionally, older adults could have answered inaccurately with “Remember” responses to compensate for memory deficits when memory-related age stereotypes were salient (Ryan & Campbell, 2021). Indeed, 29 older adults were excluded from analysis due to an insufficient number of “Know” responses. Additional work is needed to determine relations among self-reported recollection measures and the AI.

### The Autobiographical Interview is associated with subclinical depression, social cognition, and personality

From our first aim assessing the reliability and validity of the AI, we determined that, overall, the AI performed well as a measure of AM. But AM is a multifaceted construct that permeates many aspects of daily life, affecting mood, decision-making, social interactions, and personality. Given the many studies suggesting an influence of these factors on AM, our second aim was to explore how the AI associated with these related, non-mnemonic constructs. In doing so, we drew on prior observations (often in smaller samples) to inform our predictions. Leveraging our large AI sample, we sought to replicate these findings and evaluate them as precedents in the literature to guide future work.

Over-general memory has been associated with clinical depression (Brittlebank et al., 1993; Hitchcock et al., 2014; Kuyken & Dalgleish, 1995; Liu et al., 2013; Williams & Broadbent, 1986; Williams & Scott, 1988; Williams et al., 2007; Wilson & Gregory, 2018). We assessed whether AI metrics would be sensitive to this effect in sub-clinical variance of depression symptomatology. In the younger adult sample, self-reported depression was related to the recollection of more external detail across metrics. No associations were observed in older adults. This may be due to the higher recollection of external information in older adults as a group. High levels of external detail were more rare for younger adults, and may thus track symptomology. Our findings indicate that even minor depressive symptoms, below a diagnostic threshold, co-occur with less episodically rich AMs in young individuals.

The reconstructive nature of AM lends itself to a similar process by which to imagine and simulate future events (Schacter & Addis, 2007; Schacter, 2012). Indeed, a high correlation exists between internal details for past and future autobiographical events (Addis, Wong, & Schacter, 2008). This ability to envisage the future and mentally play out different scenarios, rooted in memory, has significant implications for decision-making (Boyer, 2008). For example, episodic simulation has been shown to reduce impulsive choices (Benoit et al., 2011; Peters & Büchel, 2010; Sasse et al., 2015). Such findings led to the prediction that individuals with richer episodic AM capacities are able to distance themselves from the temptation of rewards now by vividly visualizing and bringing themselves closer to their future selves (Gilbert & Wilson, 2007; Ersner-Hershfield et al., 2009; Bulley et al., 2016). Moreover, Lempert et al. (2020) reported a negative association between internal count and temporal discounting in 34 cognitively normal older adults, indicating that higher recollection of internal (specifically place, time, and perceptual) details was associated with more patient choices even into older age. In an attempt to replicate this effect in our well-powered sample, we found no associations between temporal discounting and AI metrics, including Lempert et al.’s perceptual/gist ratio. This observation warrants caution in linking internal AM to decision-making, consistent with prior reports of amnesic individuals performing within a normal range on a temporal discounting task (Kwan et al., 2015). That even semantic future thinking can minimize discounting in healthy controls (Palombo et al., 2016) further suggests that associations between future simulation and decision-making may not be specific to episodic processes.

Autobiographical recollection evokes similar patterns of brain activity as theory of mind reasoning (Buckner & Carroll, 2007). This original observation spurred much of the inquiry into understanding the relationship between AM and social cognition. For example, vivid recollection of helping another individual promotes the intention to engage in prosocial helping behavior (Gaesser & Schacter, 2014). We examined whether individual differences in empathy and theory of mind reasoning were associated with the AI metrics, predicting that more empathic individuals and those who are better at inferring emotional states would have more episodically rich memories. We observed a modest association between theory of mind, as assessed with the Reading the Mind in the Eyes task, and the AI ratio score. Scant associations were observed with trait empathy. Perhaps the induction of rich recollection or episodic simulation may be necessary to elicit prosocial behavior (Gaesser & Schacter, 2014; Gaesser & Fowler, 2020), rather than trait level covariance. The modest number of associations observed reinforce the behavioral dissociation of social and mnemonic constructs.

Given that AM revolves around the self, we sought to examine whether self-reported personality traits covaried with AI performance. Few associations were observed. Our predicted association between trait openness and internal measures did not emerge. In fact, personality did not consistently align with either internal or external measures. It could be that previous conceptions of AM (e.g., self-report, experimenter ratings) do not correspond to internal and external AI scores (see Setton et al., 2021). Notably, trait extraversion was positively associated with semantic density. More extraverted individuals are more likely to report using AM to fulfill social needs (Caci et al., 2019; Rasmussen & Berntsen, 2010; Webster, 1993), are more willing to share self-defining memories (McLean & Pasupathi, 2006), and prefer interpersonal reminiscence over other formats (Quackenbush & Barnett, 1995). Based on the social format of the AI, it is possible that more extraverted individuals share more contextual and framing details. Extraverted individuals also have greater rates of memory rehearsal, both generally and in conversation (Cappeliez & O’Rourke, 2002; Rasmussen & Berntsen, 2010; Rubin, Boals & Berntsen, 2008), suggesting that they may rehearse memories more frequently, contributing to higher rates of semantic information. Overall, correlation coefficients relating AI to the big five personality factors (DeYoung et al., 2007) were very modest. Sufficiently powered samples are needed to examine relationships between personality and AM, but are difficult to obtain given the current lengthy form of AI scoring. Advances in the scoring procedure that draw on machine learning (e.g., Peters et al., 2017; van Genugten & Schacter, 2022) will be fruitful for this purpose.

The observed associations with non-AI measures were in line with expected effect sizes for individual differences work (Gignac & Szodorai, 2016; Hemphill, 2003). However, given the number of correlations we report here, caution should be used in the interpretation of these findings. Our goal was to evaluate previously reported AM relationships with a broad range of domains, making use of a large sample of AI data. Our observations provide a stable foundation for future investigations to build upon, but additional replication is needed.

### Limited evidence for a two-factor structure in the Autobiographical Interview

In the preceding discussion, we assumed the validity of dissociating internal from external detail categories. The AI was designed to quantify and dissociate episodic from the non-episodic and semantic features of AM recall. An explicit test of the two-factor model with a CFA yielded poor model fit. Neither count nor density models outperformed a null model, and both showed large differences in residuals between the hypothesized and observed models. The CFA with density scores successfully dissociated internal and external components, but its structure was not robust. This suggests that there are nuanced relationships between sub-categories that the internal/external structure of the AI does not capture. Poor model performance may partially stem from the inclusion of semantic, repetition, and other details which demonstrated particularly low regression coefficients in the CFA with density. This is supported in theory because semantic, repetition, and other detail categories are the only external sub-categories that do not describe a spatiotemporally specific event. Indeed, given their non-mnemonic content, we anticipated that repetition and other details may not be appropriately modeled with external details. To evaluate this possibility, we conducted a CFA on internal and external details excluding the repetition and other categories. The adjusted model also fell below acceptable model thresholds. Given poor performance from our a-priori models, we conducted EFA to identify alternative AI factor structures.

Across all exploratory analyses, AI sub-categories largely separated into internal and external factors with the consistent exception of semantic, repetition, and other details. This pattern suggests that there is a robust division between external details that describe a specific event (i.e. event, place, time, perceptual, and emotion/thought details) and those that do not (i.e. semantic, repetition, and other details). Accordingly, a more granular scoring of external details may better characterize their multifaceted contents. Below we emphasize results of the full sample, which provide the highest sub-category to observation ratio. Of note, however, the dimensionality of the AI is lower in older adults, relative to the young, in both count and density analyses (see Supplemental Figures 14 and 15 and Supplemental Tables 1-4). The dimensionality of AM, and age-related changes, would be a valuable focus of future work.

The full-sample count analysis demonstrated a cross-loading of semantic, repetition, and other details between internal and external factors. In contrast, the full-sample density analysis resulted in two additional factors that covaried with (1) semantic, repetition, and other details and (2) semantic details. Internal event and perceptual details negatively loaded onto the semantic factor, again demonstrating a decoupling of episodic from semantic information that is only evident when using density measures. Internal details were also further subdivided in the full sample density analysis: internal place, time, and event details loaded onto one factor while perceptual and emotion/thought details did not. We cautiously speculate that these sub-categories could represent more concrete, verifiable details, in contrast to more subjective and perspective-driven emotion/thought and perceptual details, but further research is required to solidify this concept. Overall, these EFA findings support our supposition that density measures more readily dissociate divergent AI information. Our present results, in conjunction with novel scoring procedures (e.g., Strikwerda-Brown et al., 2019; Renoult et al., 2020), provide a substantial foundation for exploring alternative approaches to AI data.

Based on the limitations of the internal/external model, we recommend that future work consider examining sub-categories in addition to overall internal and external variables when feasible. Future evaluation of alternative models to the two-factor AI, potentially including separation of semantic, repetition, and other categories from the external category, may be worthwhile. The EFA’s suggest that alternate scoring based upon the factor structure may yield novel results. However, such an approach would require further psychometric evaluation and replication. It is inconclusive whether the poor model fit informs ongoing debate regarding the dissociation or inter-relation of episodic and semantic memory systems (e.g., Irish & Piguet, 2013; Renoult et al., 2019). However, our data strongly suggest that internal and external count scores should not be framed as distinct approximations of episodic and semantic memory given their interrelation. Instead, measures controlling for verbosity, such as density, may provide better estimates of dissociable AM components.

### Verbosity impacts the psychometric properties of the Autobiographical Interview

Across the psychometric assessment, the AI was broadly found to be a reliable and valid measure of AM. However, notable differences emerged as a function of scoring. Original count metrics for internal and external details were highly interrelated, with a shared dependency driven by the number of words spoken. These measures were also significantly linked to memory recency, as well as self-reported rehearsal.

Recently introduced density scores provide several strengths. Density scores significantly dissociate internal from external details, both in simple associations between AI metrics and the multivariate factor analyses. The internal/external distinction is essential for quantifying episodic AM. Density measures were similar across memory age, unrelated to memory rehearsal, and demonstrated larger effect sizes when comparing younger and older adults. In addition to being associated with episodic memory, internal density was also modestly correlated with executive function. External density also uniquely demonstrated negative associations with episodic memory, and positive associations with time integration (a measure of global contextual integration). Together, these observations suggest that density estimates may be an appropriate metric for individual differences in AM. However, future work is needed to investigate the greater dimensionality of the AI, based upon the AI sub-category density EFA.

## Conclusion

Publication of the AI (Levine et al., 2002) was ground-breaking, offering a novel way for researchers to distill a naturalistic, qualitative measure of memory into discrete quantities of episodic and semantic recollection. Although 20 years have passed before a comprehensive psychometric assessment, the AI is a highly reliable and valid measure of AM. By deriving and testing several AI metrics, here we find that verbosity-controlled variables may be better suited for detecting individual differences that reflect recollective processes beyond narrative style. The study of trait-level AM has fostered a deeper understanding of how AM and non-mnemonic cognition interact. It has also moved the field toward uncovering how individual variation in brain structure and function shape the way in which humans recall the personal past, at scales ranging from the biophysical properties of axons (Clark et al., 2022) resting-state functional connectivity (Setton et al., 2022a), and temporal lobe volumetry (Setton et al., 2022b). However, several open questions remain, such as the appropriate factor structure for the AI and how it covaries with other related domains such as perception, language, and attention. Updates to scoring protocols that consider alternate models and expedite the scoring process will be instrumental for filling in these gaps to systematically assess AM in ever larger samples.

## Acknowledgements

This project was supported by the Canadian Institute of Health Research and the Natural Sciences and Engineering Research Council of Canada. R.N.S. is a Research Scholar supported by the Fonds de la Recherche du Quebec - Santé.

## CRediT author statement

**Amber W. Lockrow:** Conceptualization, Methodology, Formal analysis, Investigation, Data Curation, Writing - Original Draft, Review & Editing, Visualization; **Roni Setton:** Data Curation, Writing - Review & Editing; **Karen A.P. Spreng**: Resources, Writing - Review & Editing; **Signy Sheldon**: Writing - Review & Editing; **Gary R. Turner:** Writing - Review & Editing, Supervision, Project administration, Funding acquisition; **R. Nathan Spreng:** Conceptualization, Methodology, Writing - Original Draft, Review & Editing, Supervision, Project administration, Funding acquisition

## Data availability

The dataset generated and analysed for the current study are available through the Open Science Framework, project “Psychometrics of Autobiographical Memory” contributed by R.N.S. (https://osf.io/fzkm7/) or from the corresponding author. The authors have no relevant financial or non-financial interests to disclose.

## Supplemental Materials

**Supplemental Figure 1.**
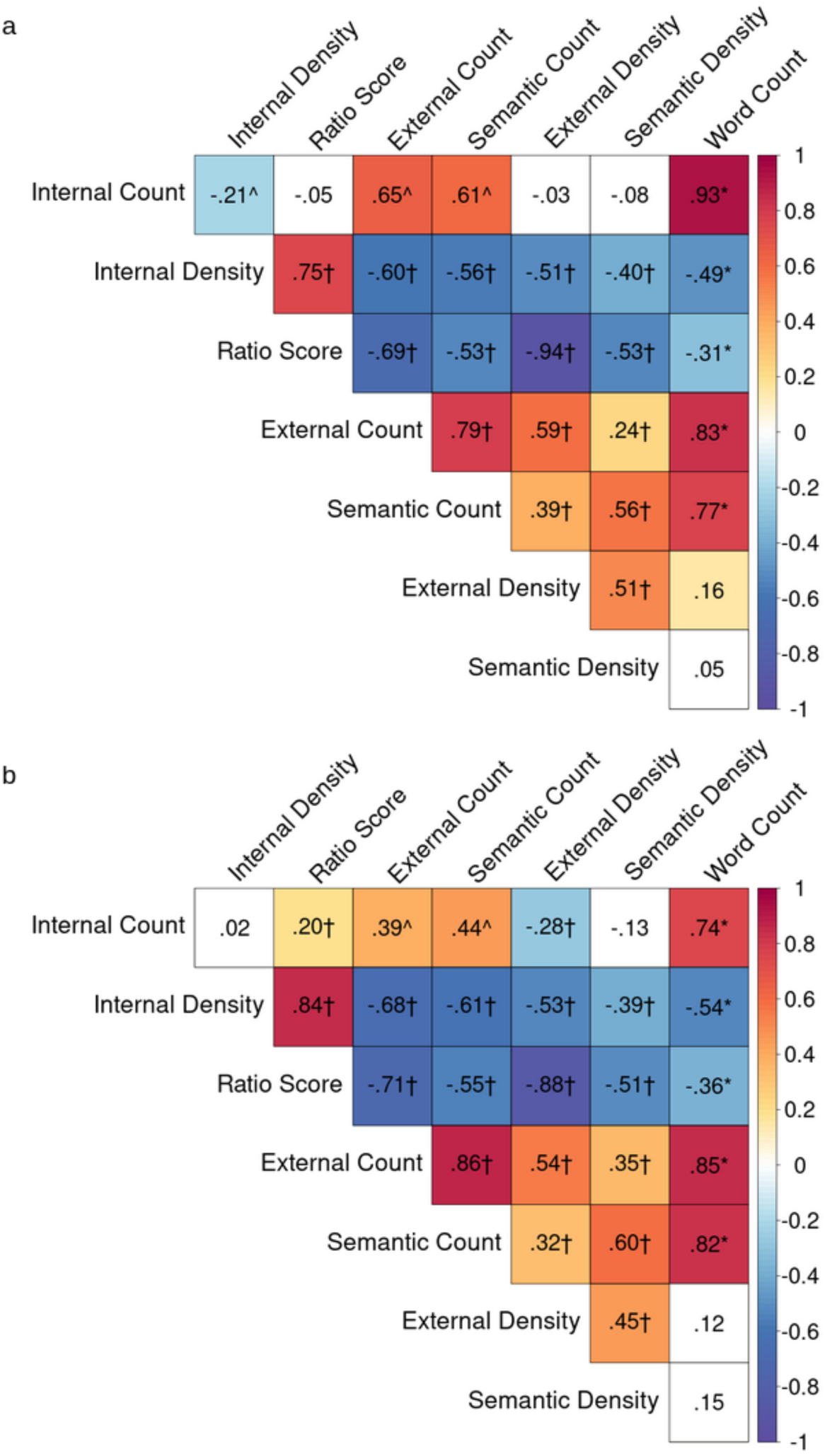
Internal Consistency of the Autobiographical Interview: Measures of Interest in Younger and Older Adults. Partial product-moment correlations controlling for gender between the 8 AI measures of interest in a) younger adults only b) older adults only. The matrix contains partial r values, which are color-coded by strength of the correlation. Cells with a white background indicate non-significant associations (p > .05, uncorrected). Color cells are significant (p < .05). † significant predicted association, * unpredicted significant association with Bonferroni correction, ^ associations opposite to prediction (p < .05, uncorrected).

**Supplemental Figure 2.**
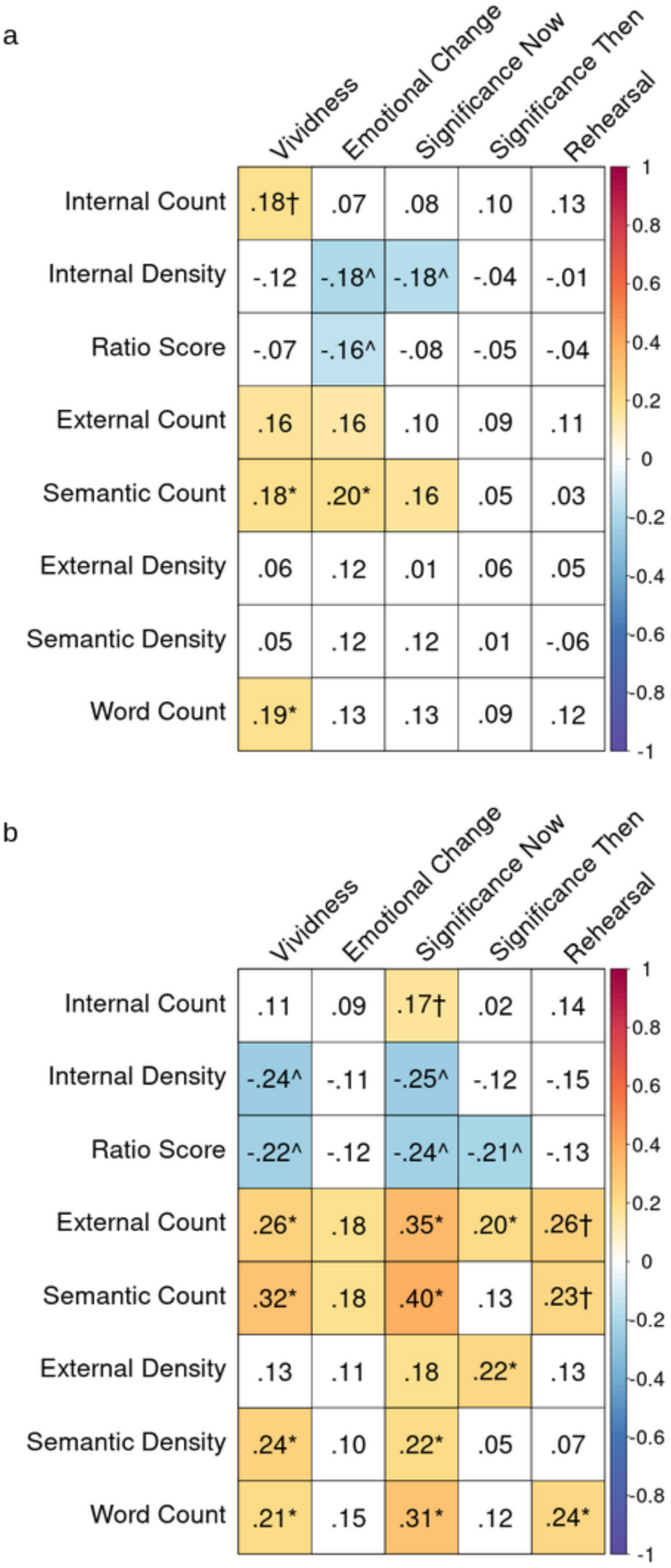
Internal Consistency of the Autobiographical Interview: Participant Self-Report Ratings in Younger and Older Adults. Partial Spearman correlations controlling for gender between the 8 AI measures of interest and self-report ratings controlling for gender in a) younger adults and b) older adults. The matrix contains partial ρ values, which are color-coded by strength of the correlation. Cells with a white background indicate non-significant associations (p > .05, uncorrected). Color cells are significant (p < .05). † significant predicted association, * unpredicted significant association with Bonferroni correction, ^ associations opposite to prediction (p < .05, uncorrected)

**Supplemental Figure 3.**
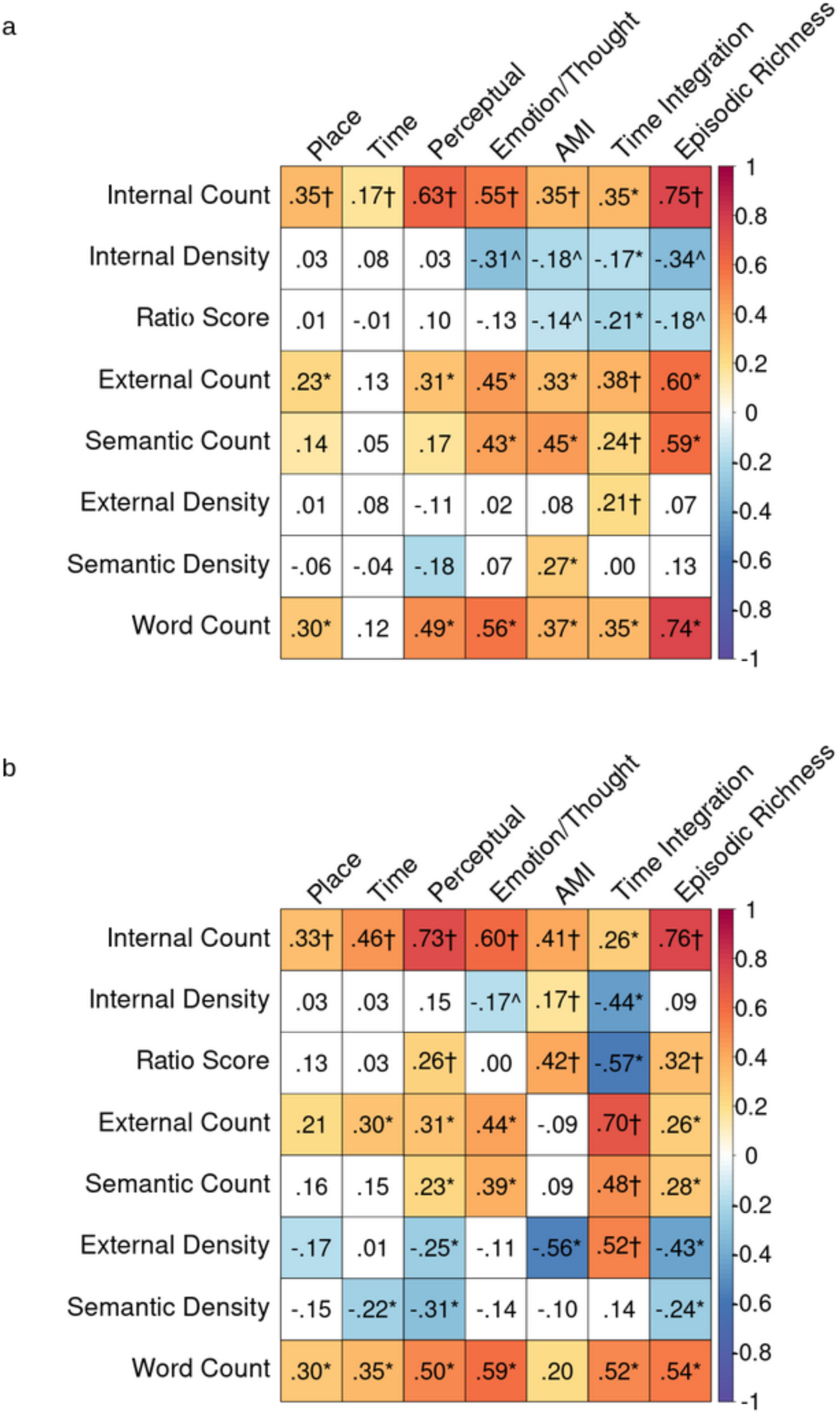
Internal Consistency of the Autobiographical Interview: Scorer Ratings in Younger and Older Adults. AMI=episodic specificity rating from the Autobiographical Incident Schedule of the Autobiographical Memory Interview. Partial Spearman Correlations between the 8 AI measures of interest and scorer overall ratings controlling for gender in a) younger adults only b) older adults only. The matrix contains partial ρ values, which are color-coded by strength of the correlation. Cells with a white background indicate non-significant associations (p > .05, uncorrected). Color cells are significant (p < .05). † significant predicted association, * unpredicted significant association with Bonferroni correction, ^ associations opposite to prediction (p < .05, uncorrected).

**Supplemental Figure 4.**
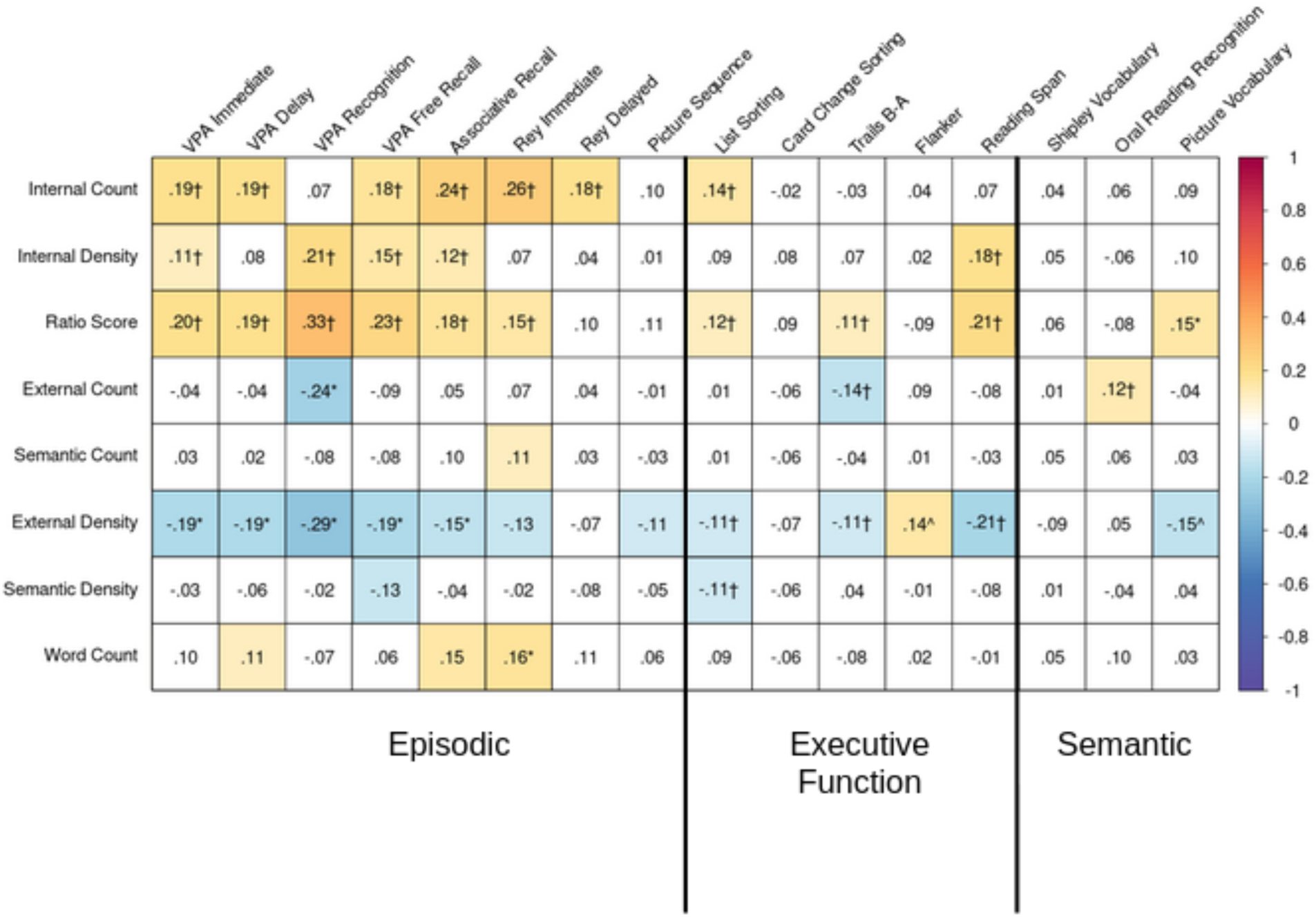
Convergent Validity of the Autobiographical Interview with Episodic Memory, Executive Function, and Semantic Memory Index Scores. Partial product-moment correlations controlling for age and gender across all participants between the 8 AI measures of interest and index scores in all participants. The matrix contains partial r values, which are color-coded by strength of the correlation. Cells with a white background indicate non-significant associations (p > .05, uncorrected). Color cells are significant (p < .05). † significant predicted association, * unpredicted significant association with Bonferroni correction.

**Supplemental Figure 5.**
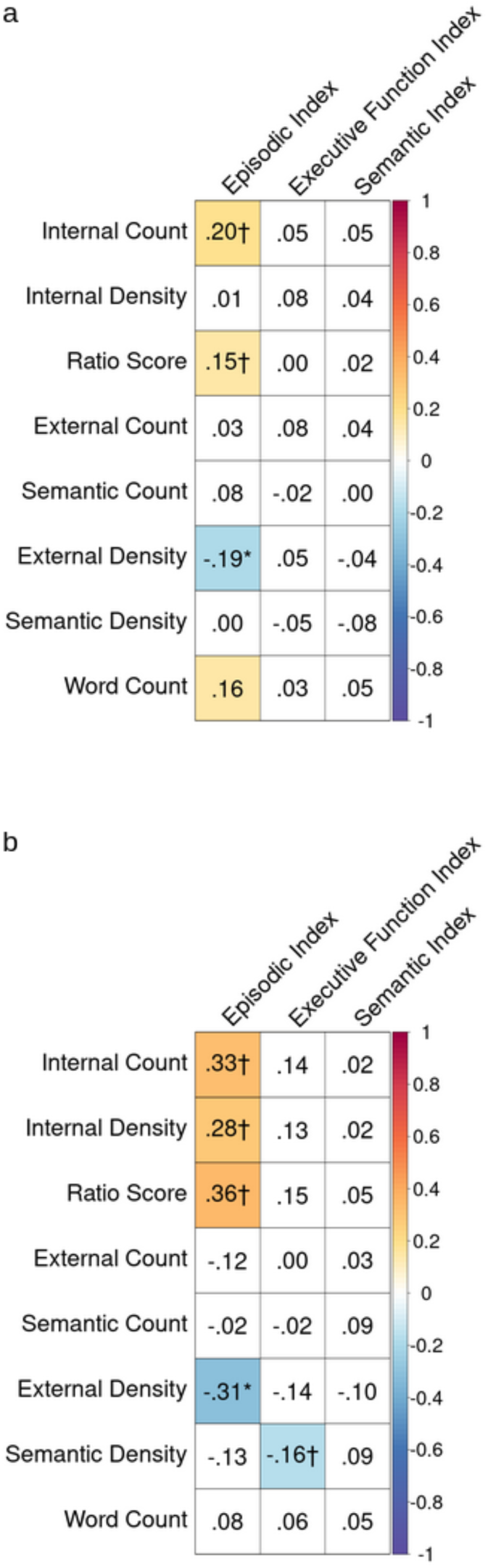
Convergent Validity of the Autobiographical Interview with Episodic Memory, Executive Function, and Semantic Memory Index Scores in Younger and Older Adults. Partial product-moment correlations controlling for gender between index scores and the 8 AI measures of interest in a) younger adults and b) older adults. The matrix contains partial r values, which are color-coded by strength of the correlation. Cells with a white background indicate non-significant associations (p > .05, uncorrected). Color cells are significant (p < .05). † significant predicted association, * unpredicted significant association with Bonferroni correction.

**Supplemental Figure 6.**
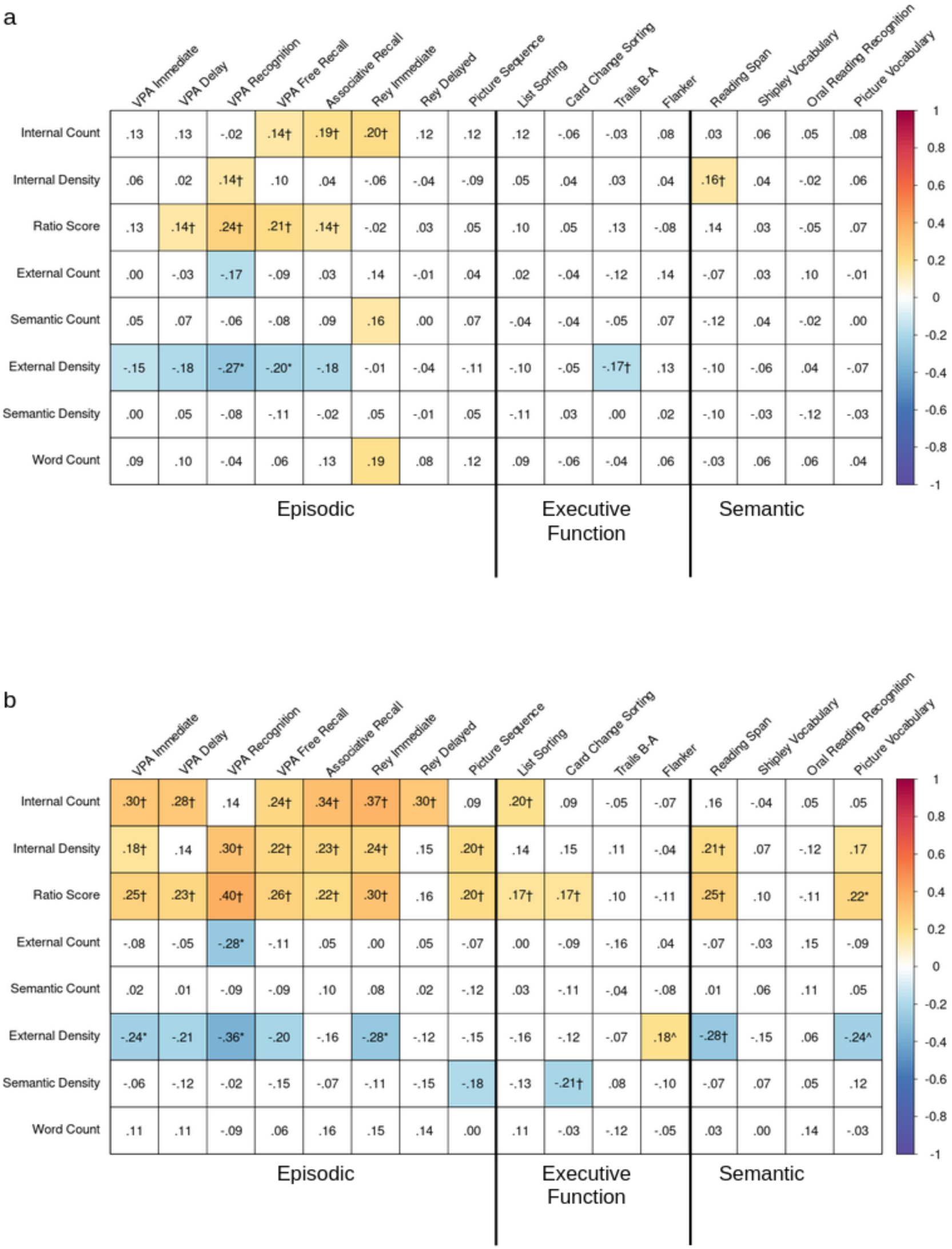
Convergent Validity of the Autobiographical Interview with Episodic Memory, Executive Function, and Semantic Memory Index Scores in Younger and Older Adults. Partial product-moment correlations controlling for gender between the 8 AI measures of interest and index scores in a) younger adults and b) older adults. The matrix contains partial r values, which are color-coded by strength of the correlation. Cells with a white background indicate non-significant associations (p > .05, uncorrected). Color cells are significant (p < .05). † significant predicted association, * unpredicted significant association with Bonferroni correction, ^ associations opposite to prediction (p < .05, uncorrected)

**Supplemental Figure 7.**
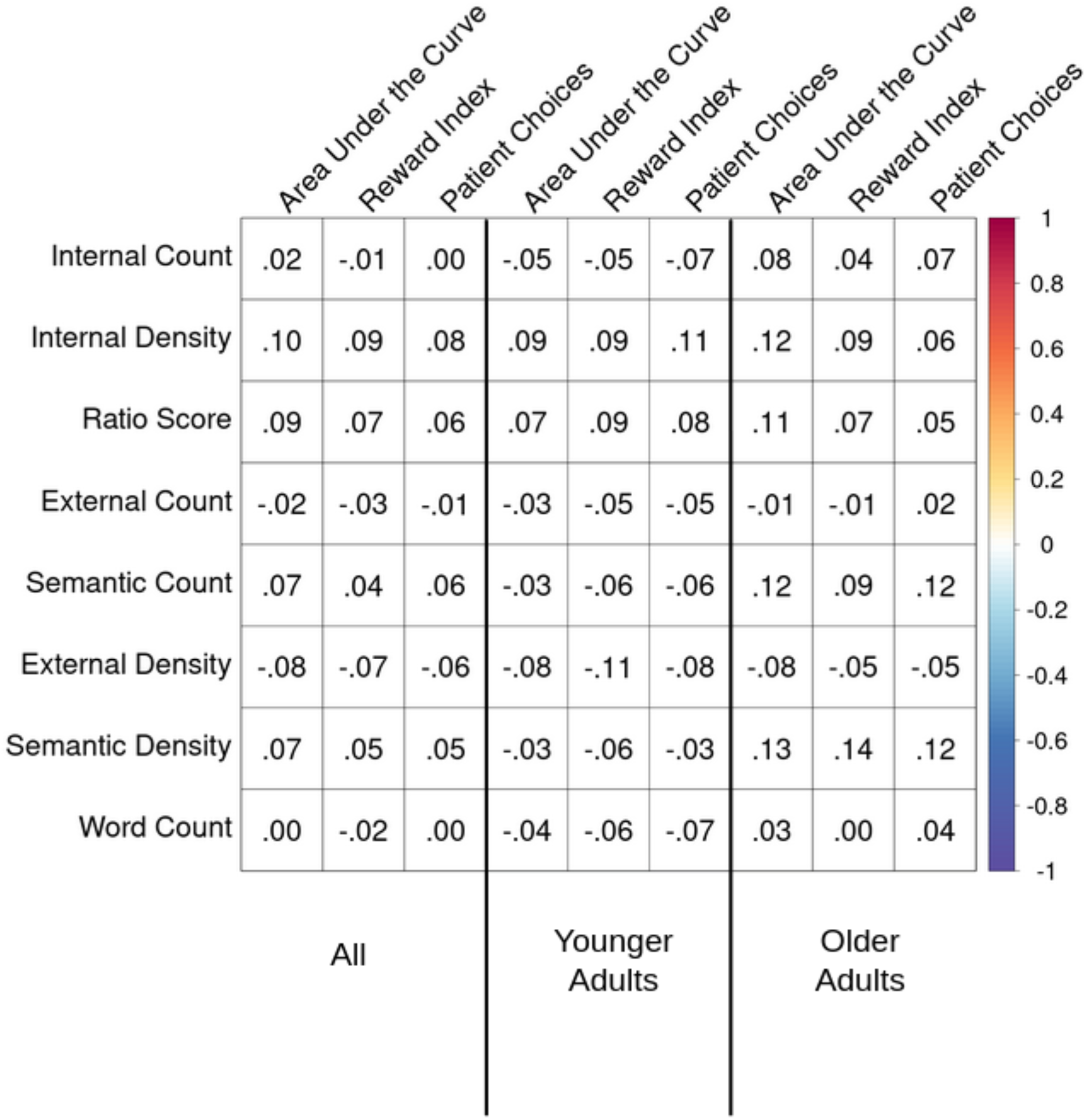
Associations between the Autobiographical Interview with Measures of Temporal Discounting. Partial product-moment correlations between the 8 AI measures of interest and measures of temporal discounting across all participants controlling for age and gender and within each age group controlling for gender. The matrix contains partial r values, which are color-coded by strength of the correlation. Cells with a white background indicate no significant association (p > .05, uncorrected)

**Supplemental Figure 8.**
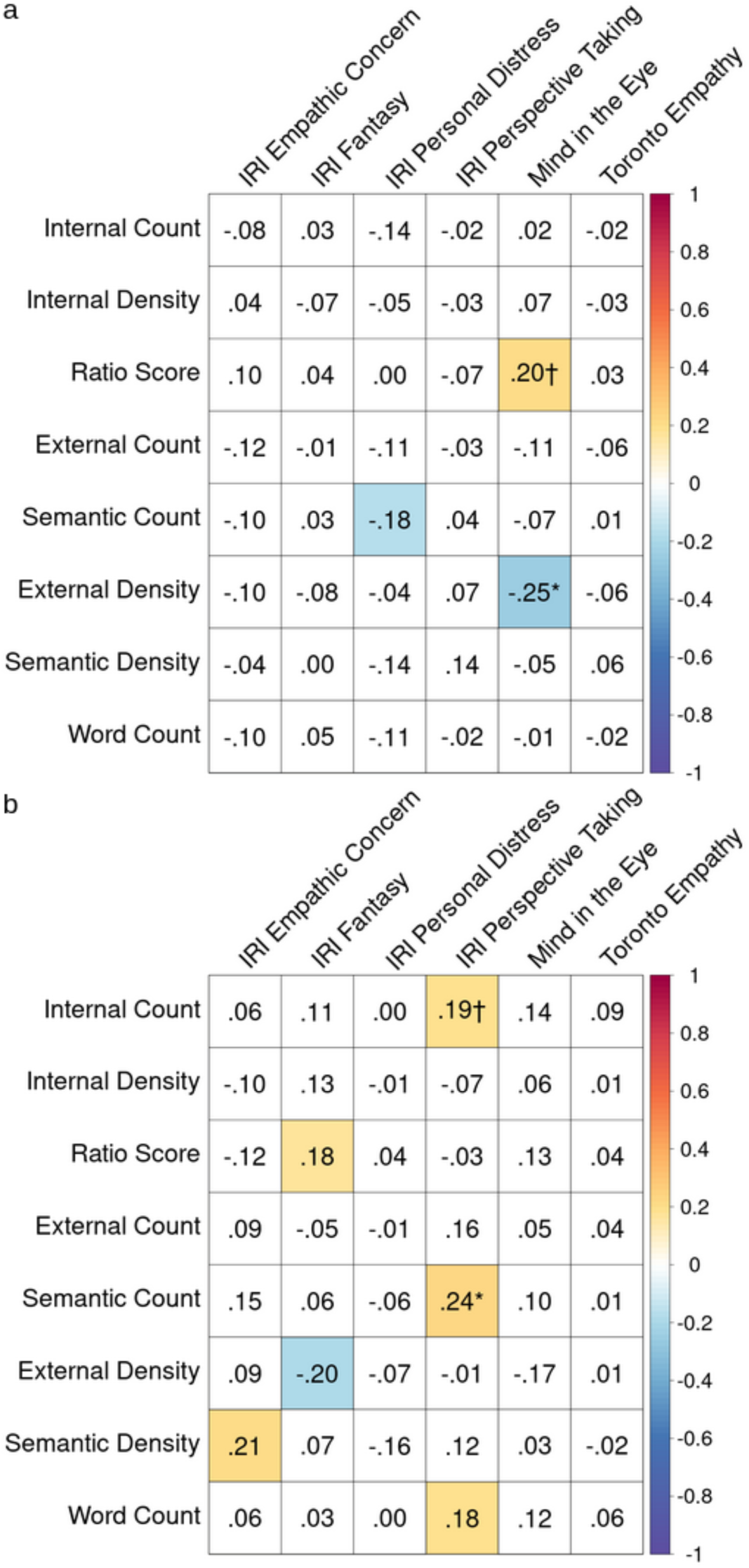
Convergent Validity of the Autobiographical Interview with Measures of Social Cognition in Younger and Older Adults. IRI=Interpersonal Reactivity Index. Partial product-moment correlations between the 8 AI measures of interest and measures of personality controlling for gender within a) younger adults and b) older adults. The matrix contains partial r values, which are color-coded by strength of the correlation. Cells with a white background indicate non-significant associations (p > .05, uncorrected). Color cells are significant (p < .05). † significant predicted association, * unpredicted significant association with Bonferroni correction.

**Supplemental Figure 9.**
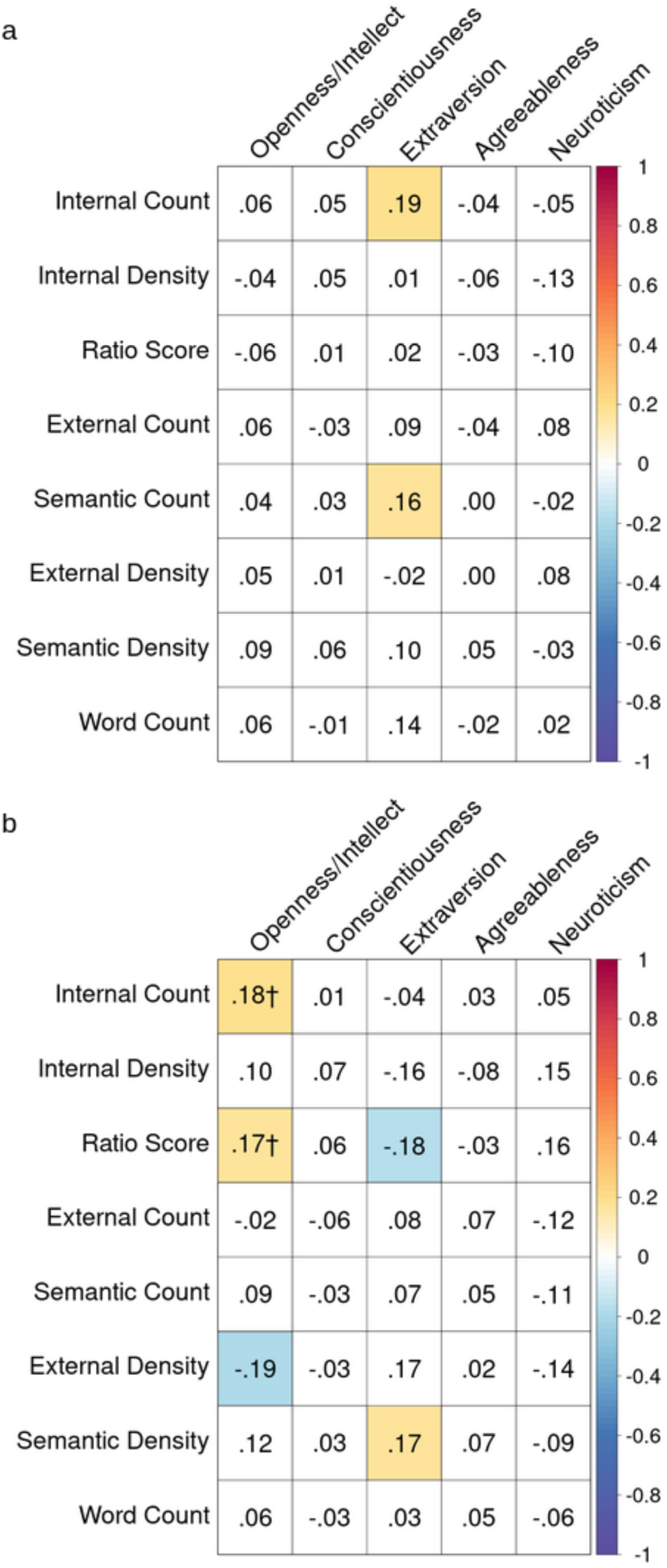
Associations between the Autobiographical Interview with Measures of Personality in Younger and Older Adults. Partial product-moment correlations between the 8 AI measures of interest and measures of personality controlling for gender within a) younger adults and b) older adults. The matrix contains partial r values, which are color-coded by strength of the correlation. Cells with a white background indicate non-significant associations (p > .05, uncorrected). Color cells are significant (p < .05). † significant predicted associat

**Supplemental Figure 10.**
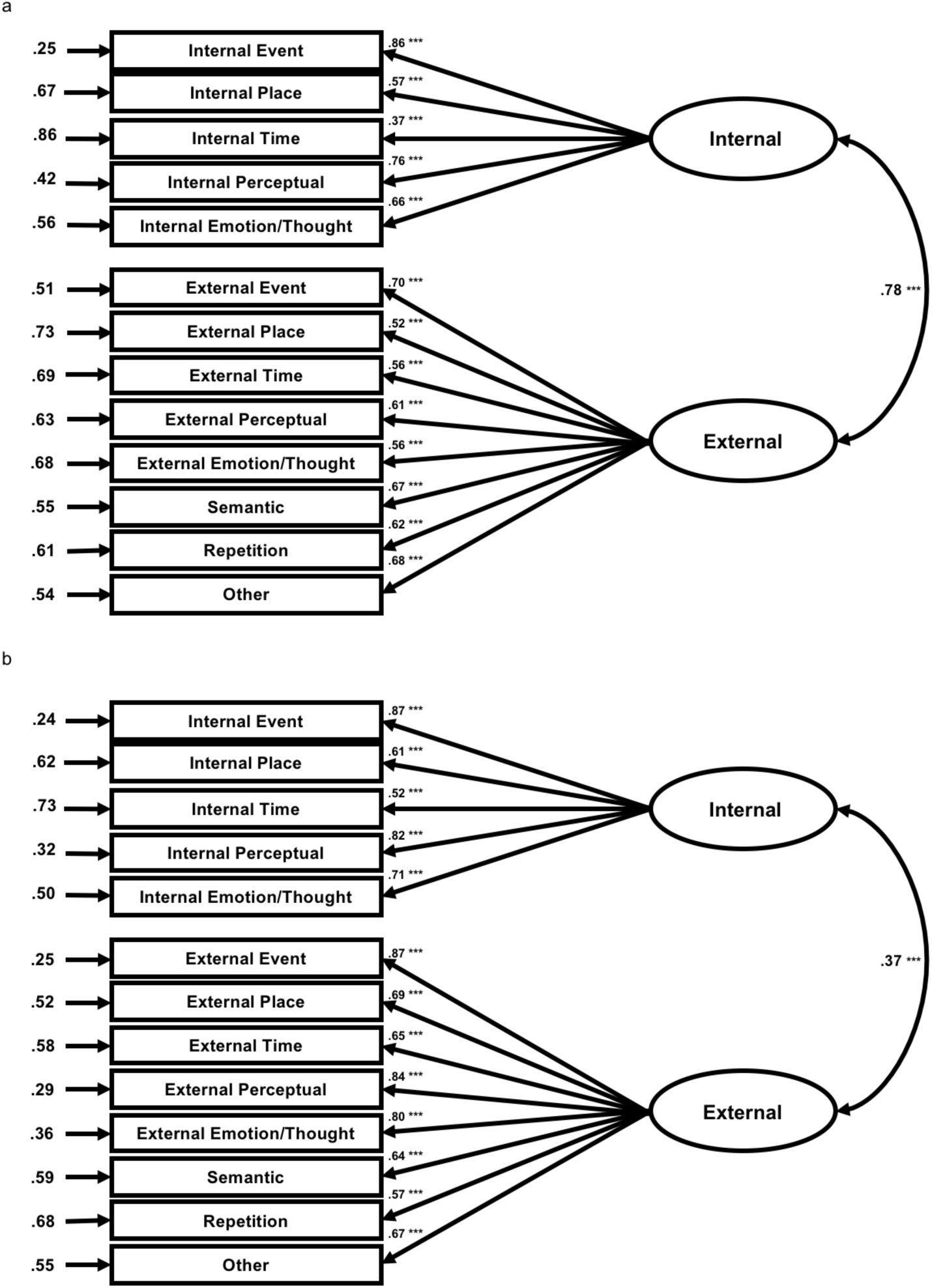
Confirmatory Factor Analysis of Internal and External Detail Counts. The ellipses represent the latent variables of internal and external counts whereas the rectangles represent observed sub-categories of detail types which fall into each latent variable category, as represented by the straight arrows. Straight arrows between the latent and observed variables are shown with adjacent standardized regression coefficients (beta weights) and estimated with maximum likelihood estimation. The curved arrow represents the correlational relationship between the two latent variables and is shown with its adjacent correlation coefficient. Numbers at the end of the shorter arrows are standardized error variances. Diagrams are shown separately for a) younger adults and b) older adults, with groups analyzed in the same model. n.s. not significant; *p<.05; **p < .01; ***p<.001.

**Supplemental Figure 11.**
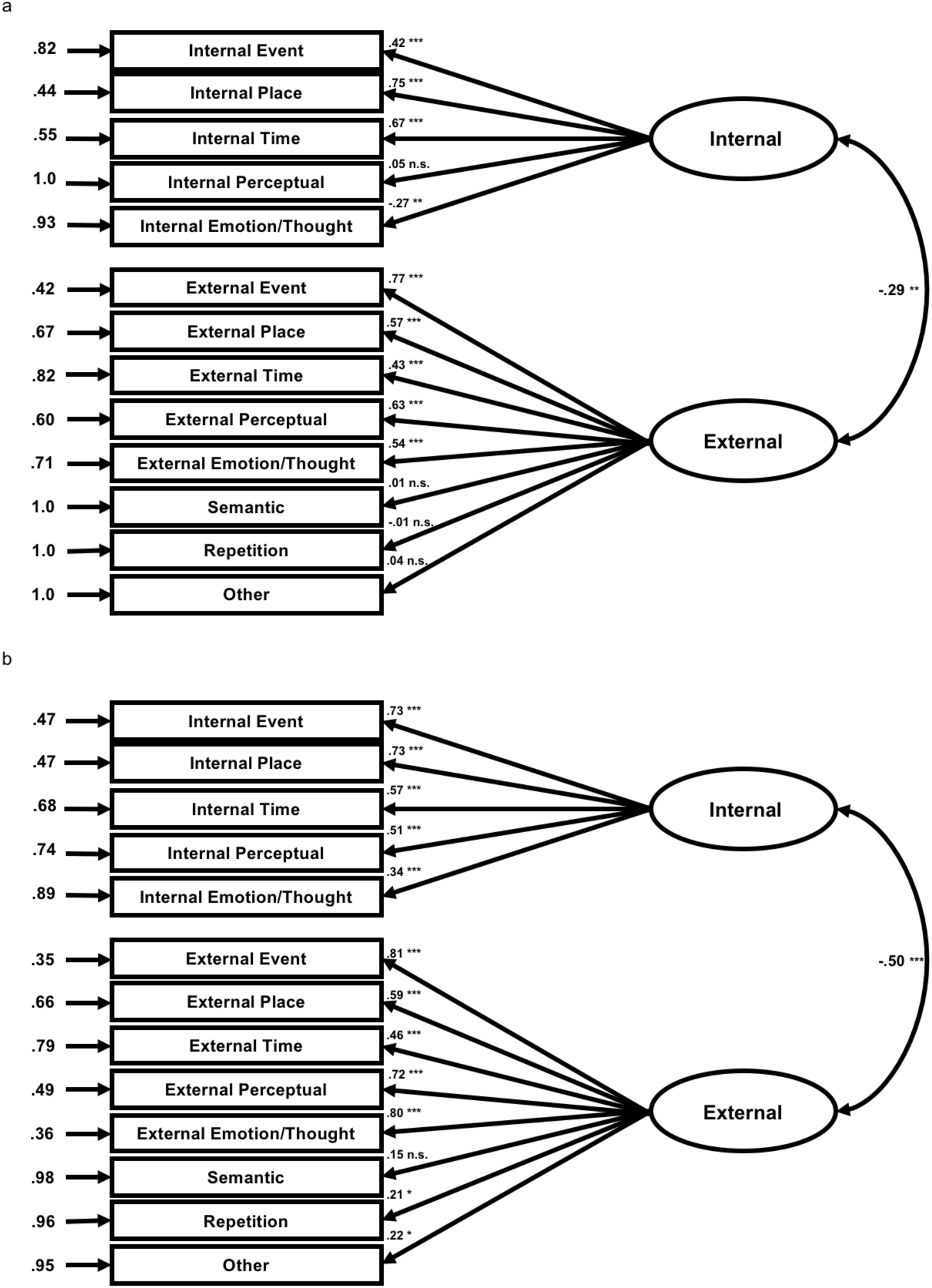
Confirmatory Factor Analysis of Internal and External Density Scores. n.s. not significant; Diagrams are shown separately for a) younger adults and b) older adults, with groups analyzed in the same model. n.s. not significant; *p<.05; **p < .01; ***p<.001.

**Supplemental Figure 12.**
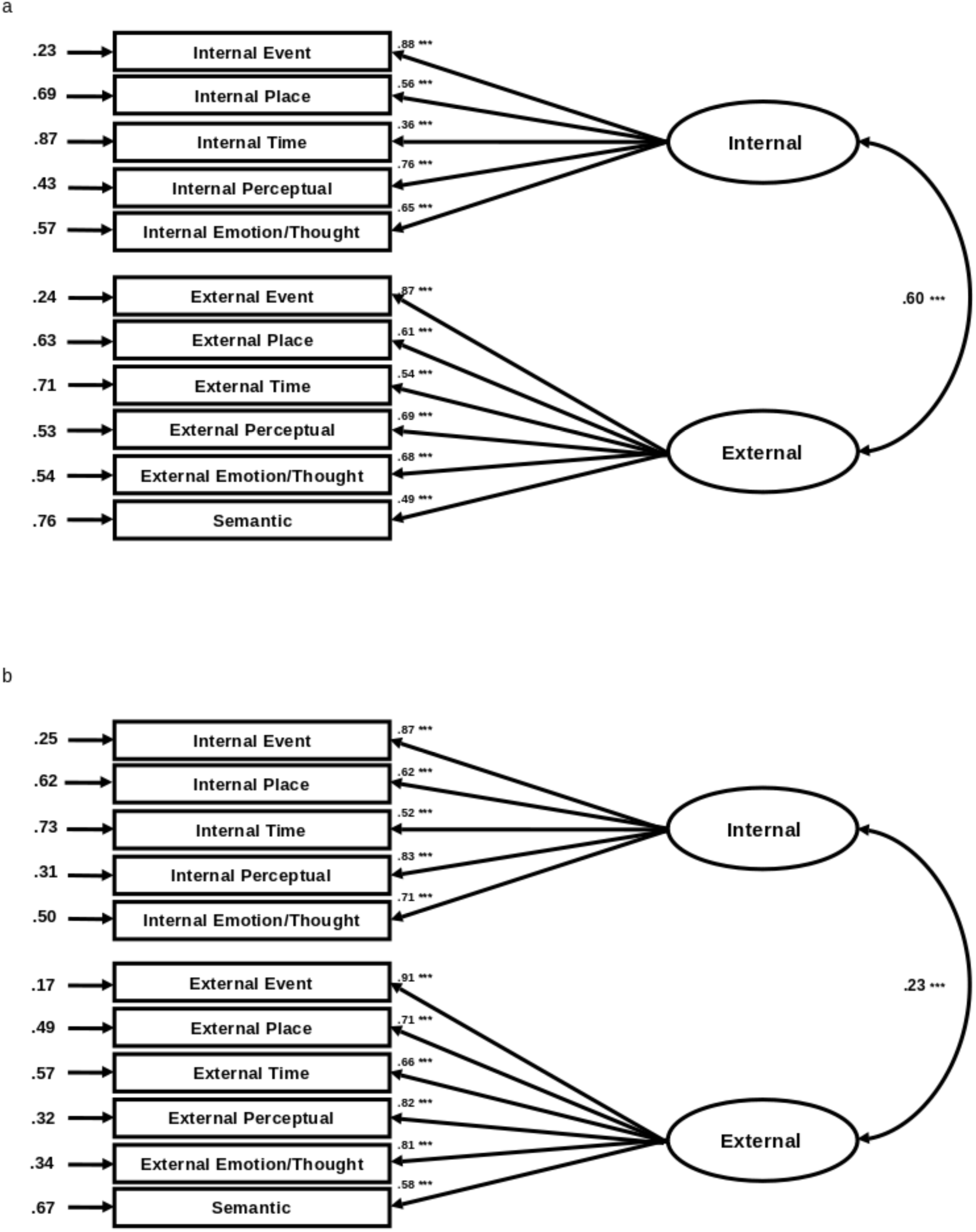
Confirmatory Factor Analysis of Internal and External Detail Counts Excluding Repetition and Other Details. N.s. not significant; Diagrams are shown separately for a) younger adults and b) older adults, with groups analyzed in the same model. N.s. not significant; *p<.05; **p < .01; ***p<.001.

**Supplemental Figure 13.**
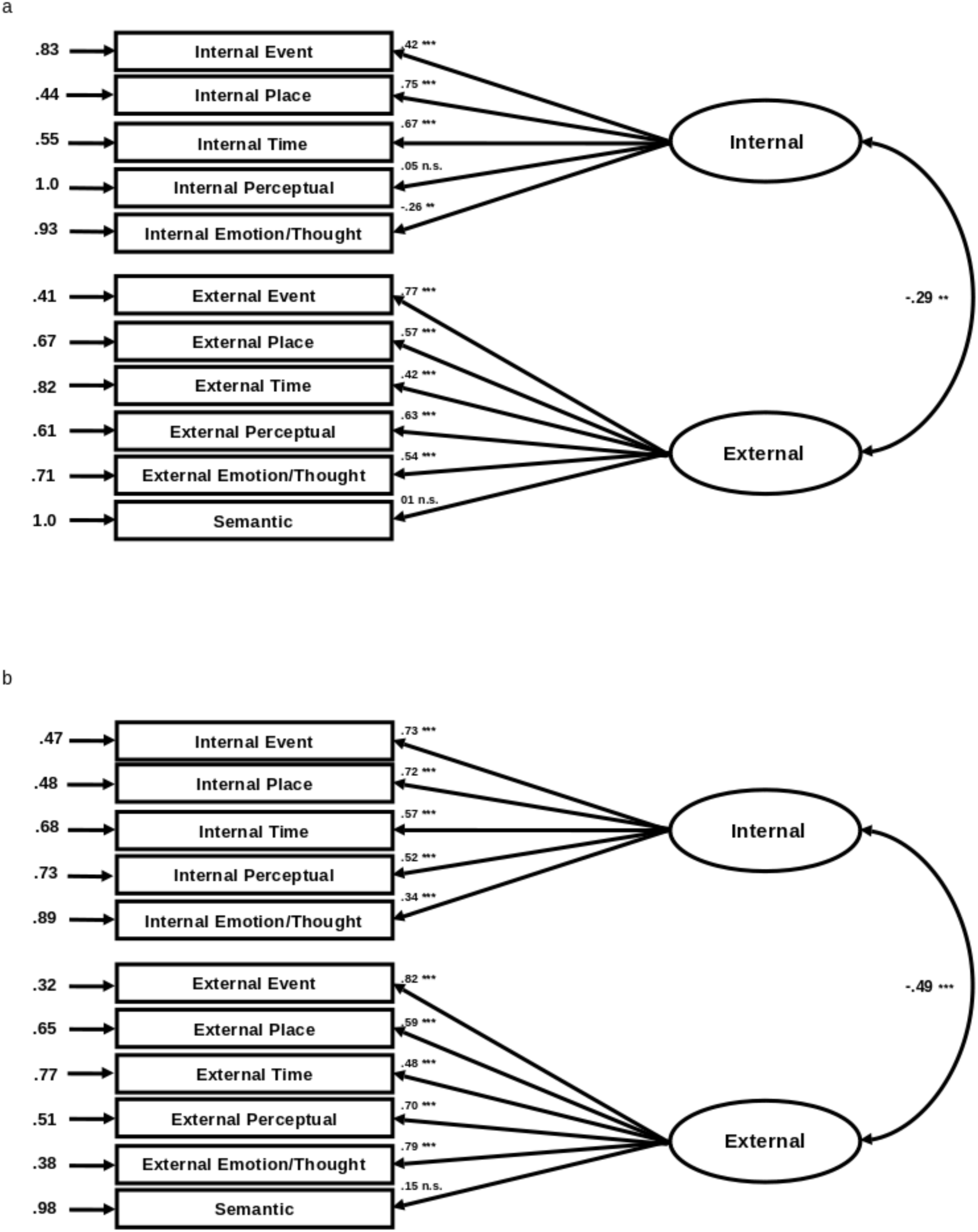
Confirmatory Factor Analysis of Internal and External Density Scores Excluding Repetition and Other Details. n.s. not significant; Diagrams are shown separately for a) younger adults and b) older adults, with groups analyzed in the same model. n.s. not significant; *p<.05; **p < .01; ***p<.001.

**Supplemental Figure 14.**
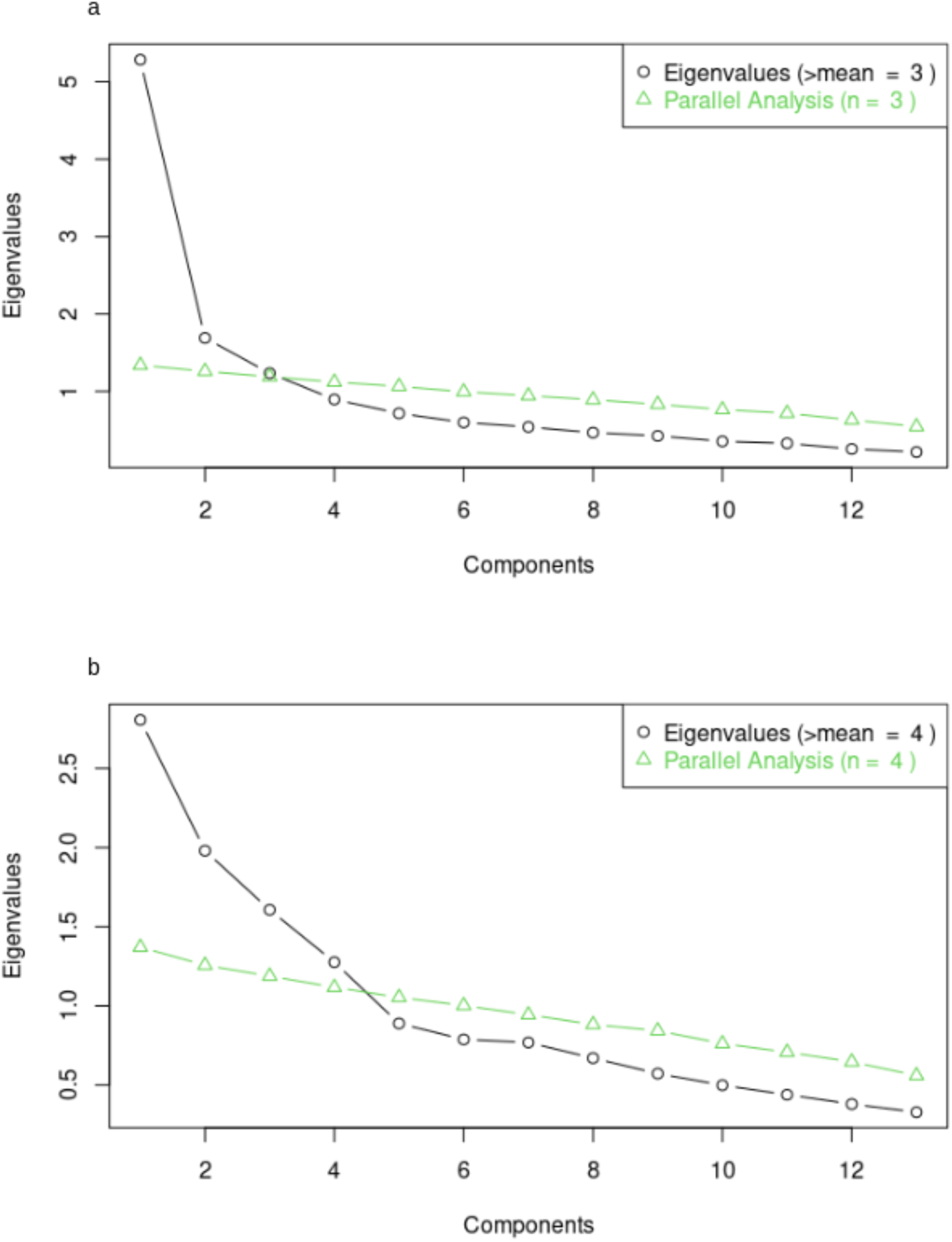
Exploratory Factor Analysis of Autobiographical Interview Detail Sub-categories in Younger Adults. Resulting eigenvalues and parallel analysis results for an optimal number of factors from an exploratory factor analysis of a) internal and external details and b) internal and external density scores.

**Supplemental Table 1.** Factor Loadings for Exploratory Factor Analysis of Detail Sub-category Count in Younger Adults. Factor loadings from exploratory factor analysis with varimax rotation for each of three factors onto sub detail types. Factor columns are sorted by variance explained (most to least). Loadings greater than .3 are bolded.

| Detail type | Factor loading |  |  |
| --- | --- | --- | --- |
|  | 1 | 2 | 3 |
| Internal Emotional Details | **.75** | .22 | -.02 |
| Internal Event Details | **.73** | .24 | .25 |
| Internal Perceptual Details | **.64** | .11 | **.33** |
| External Semantic Details | **.61** | .25 | .21 |
| External Other Details | **.57** | .17 | **.44** |
| External Repetitions | **.56** | .10 | **.40** |
| Internal Place Details | **.36** | .04 | **.58** |
| External Emotional Details | **.32** | **.68** | -.12 |
| External Event Details | .27 | **.84** | .10 |
| External Perceptual Details | .16 | **.65** | .28 |
| External Place Details | .04 | **.58** | .27 |
| External Time Details | .16 | **.41** | **.47** |
| Internal Time Details | .13 | .15 | **.51** |

**Supplemental Table 2.** Factor Loadings for Exploratory Factor Analysis of Detail Sub-category Density Scores in Younger Adults. Factor loadings from exploratory factor analysis with varimax rotation for each of four factors onto sub detail types. Factor columns are sorted by variance explained (most to least). Loadings greater than .3 or less than -.3 are bolded.

| Detail type | Factor loading |  |  |  |
| --- | --- | --- | --- | --- |
|  | 1 | 2 | 3 | 4 |
| External Event Details | **.79** | -.24 | -.14 | .12 |
| External Perceptual Details | **.66** | .17 | -.05 | -.09 |
| External Place Details | **.58** | .14 | .09 | .18 |
| External Emotional Details | **.47** | -.16 | -.29 | .16 |
| External Time Details | **.46** | .23 | .10 | -.04 |
| External Other Details | .06 | **.58** | -.02 | -.02 |
| External Repetitions | -.02 | **.56** | -.01 | .12 |
| External Semantic Details | -.07 | **.35** | -.09 | **.59** |
| Internal Place Details | -.16 | -.07 | **.70** | -.13 |
| Internal Time Details | -.03 | -.22 | **.62** | -.09 |
| Internal Event Details | -.21 | **-.65** | **.30** | -.02 |
| Internal Emotional Details | -.10 | -.03 | **-.39** | -.18 |
| Internal Perceptual Details | -.12 | .05 | -.01 | **-.48** |

**Supplemental Figure 15.**
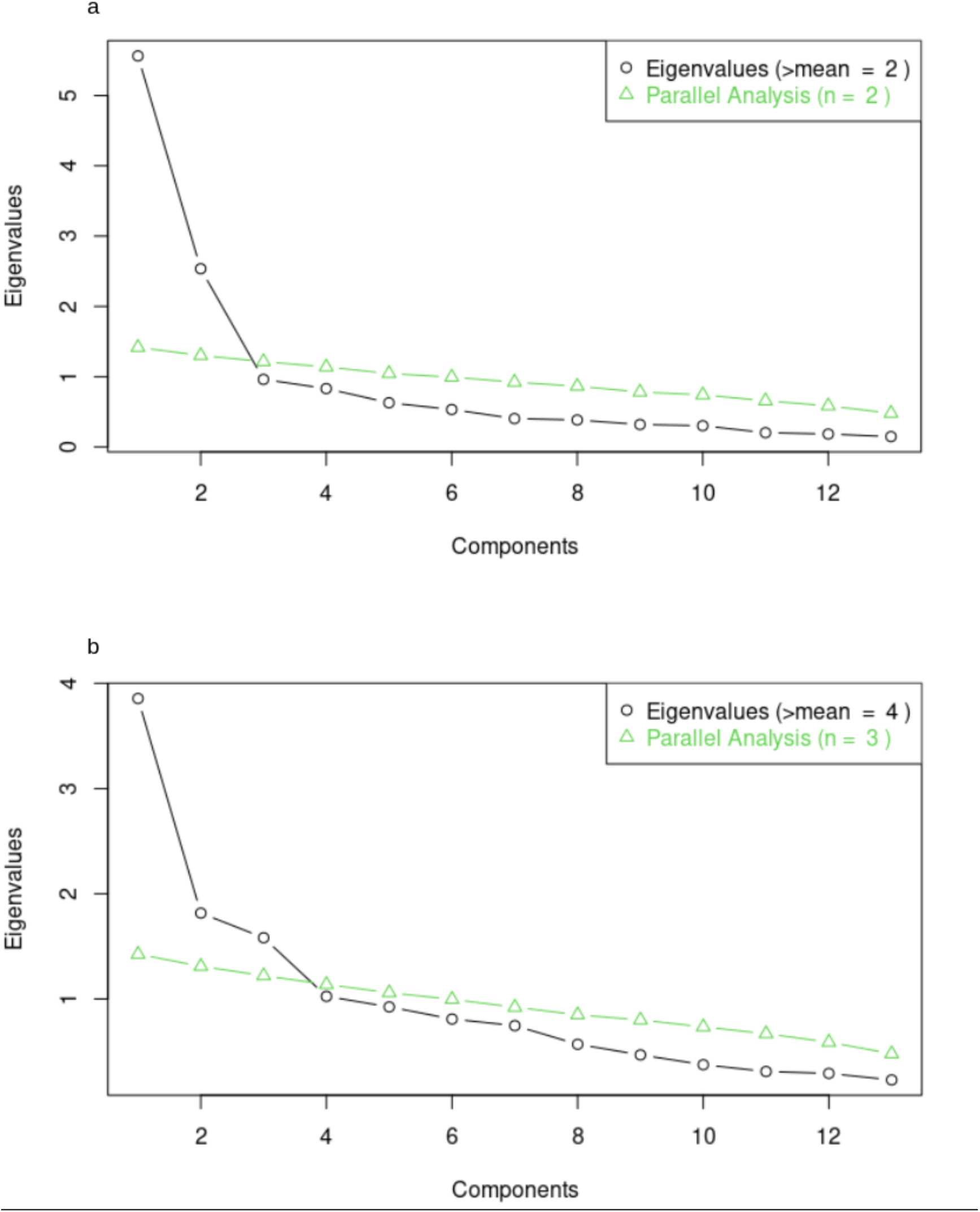
Exploratory Factor Analysis of Autobiographical Interview Detail Sub-categories in Older Adults. Resulting eigenvalues and parallel analysis results for an optimal number of factors from an exploratory factor analysis of a) internal and external details and b) internal and external density scores.

**Supplemental Table 3.** Factor Loadings for Exploratory Factor Analysis of Detail Sub-category Count in Older Adults. Factor loadings from exploratory factor analysis with varimax rotation for each of two factors onto sub detail types. Factor columns are sorted by variance explained (most to least). Loadings greater than .3 are bolded.

| Detail type | Factor loading |  |
| --- | --- | --- |
|  | 1 | 2 |
| External Event Details | **.91** | .06 |
| External Emotional Details | **.83** | .05 |
| External Perceptual Details | **.80** | .20 |
| External Place Details | **.67** | .05 |
| External Time Details | **.63** | .12 |
| External Other Details | **.57** | **.53** |
| External Semantic Details | **.54** | **.49** |
| External Repetitions | **.44** | **.55** |
| Internal Perceptual Details | -.03 | **.87** |
| Internal Event Details | .16 | **.83** |
| Internal Emotional Details | .20 | **.69** |
| Internal Place Details | -.02 | **.60** |
| Internal Time Details | .13 | **.49** |

**Supplemental Table 4.**
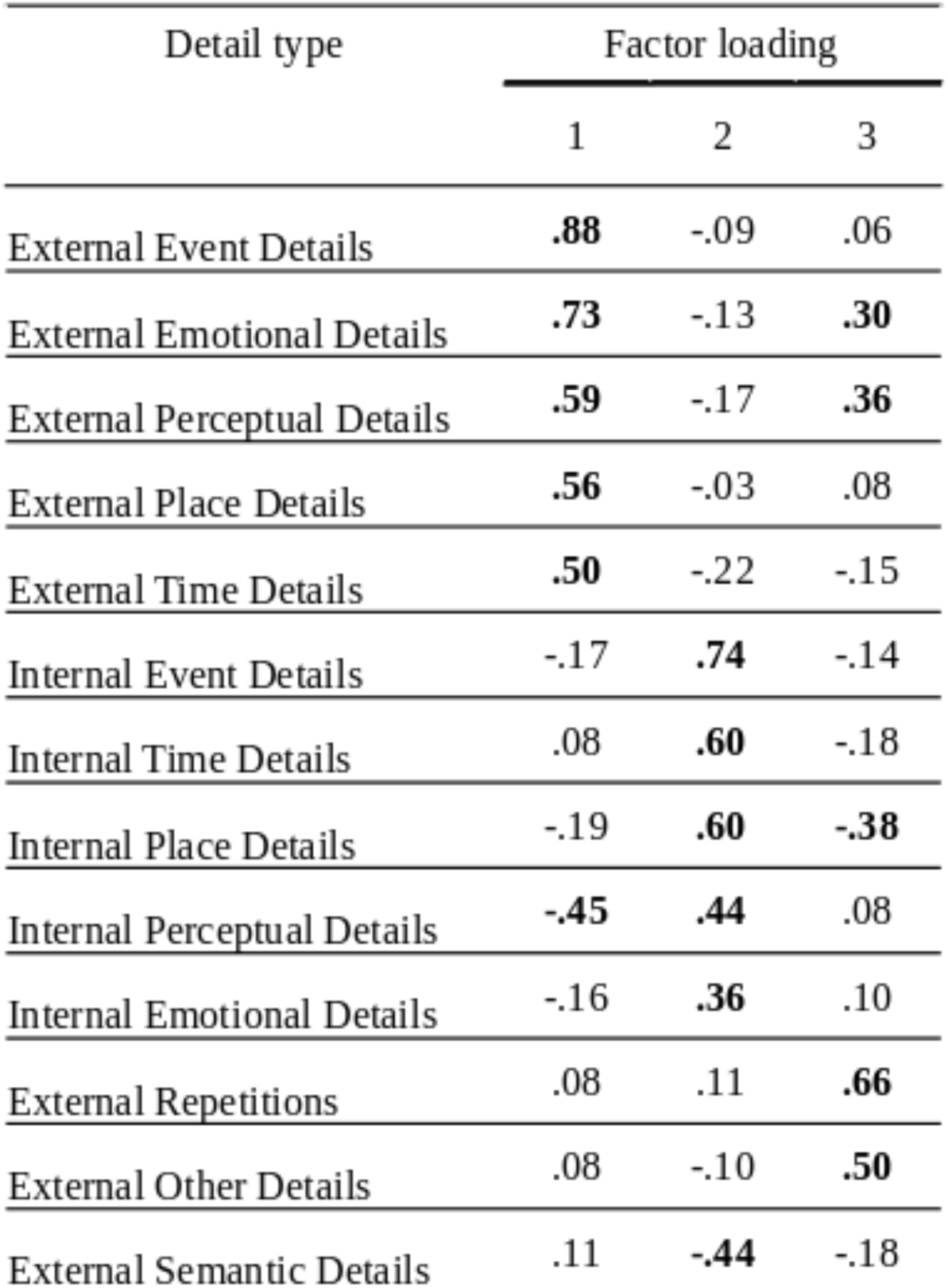
Factor Loadings for Exploratory Factor Analysis of Detail Sub-category Density Scores in Older Adults. Factor loadings from exploratory factor analysis with varimax rotation for each of three factors onto sub detail types. Factor columns are sorted by variance explained (most to least). Loadings greater than .3 or less than -.3 are bolded.

